# Arabidopsis cell suspension culture that lacks circadian rhythms can be recovered by constitutive ELF3 expression

**DOI:** 10.1101/2022.05.12.491735

**Authors:** Kanjana Laosuntisuk, Jigar S. Desai, Colleen J. Doherty

## Abstract

Callus and cell suspension culture techniques are valuable tools in plant biotechnology and are widely used in fundamental and applied research. For studies in callus and cell suspension cultures to be relevant, it is essential to know if the underlying biochemistry is similar to intact plants. This study examined the expression of core circadian genes in Arabidopsis callus from the cell suspension named AT2 and found that the circadian rhythms were impaired. The circadian waveforms are similar to intact plants in the light/dark cycles, but the circadian expression in the AT2 callus stopped in the free-running, constant light conditions. Temperature cycles could drive the rhythmic expression in constant conditions, but there were novel peaks at the point of temperature transitions unique to each clock gene. We found that callus freshly induced from seedlings had normal oscillations, like intact plants, suggesting that the loss of the circadian oscillation in the AT2 callus was specific to this callus. We determined that neither the media composition nor the source of the AT2 callus caused this disruption. We observed that *ELF3* expression was not differentially expressed between dawn and dusk in both entrained, light-dark cycles and constant light conditions. Overexpression of *ELF3* in the AT2 callus partially recovers the circadian oscillation in the AT2 callus. This work shows that while callus and cell suspension cultures can be valuable tools for investigating plant responses, careful evaluation of their phenotype is important. Moreover, the altered circadian rhythms under constant light and temperature cycles in the AT2 callus could be useful backgrounds to understand the connections driving circadian oscillators and light and temperature sensing at the cellular level.

## 1. Introduction

Research progress in plant science can be limited by the time required to grow and transform plant species. Callus and cell suspension culture systems address this limitation. Investigations into plant cellular responses can be carried out rapidly in callus and cell suspension culture systems. Callus can be transformed and mutagenized enabling reporter, overexpression, and loss of function approaches. These systems could facilitate bioengineering applications and interest in increased production of specialized metabolites (Efferth 2019). However, the functionality of callus and cell suspension culture systems depends on their ability to approximate the cellular functions of the whole plant reliably. Previously, there have been conflicting reports on the persistence of the circadian clock in callus and cell suspension cultures (Nakamichi et al. 2003, 2004). We investigated the circadian clock in an Arabidopsis cell suspension line, AT2 (Tanurdzic et al. 2008), to evaluate this system’s performance, potential, and limitations for callus and plant tissue culture-based studies.

Circadian clocks are biological mechanisms that maintain a 24-hour period regardless of external cues (McClung 2006). Circadian clocks exist in a wide range of organisms, from bacteria to mammals and plants (Jabbur, Zhao, and Johnson 2021). Robust clocks result in improved growth and fitness under environmental changes (Dodd et al. 2005; Green et al. 2002; Hotta et al. 2007; Yerushalmi and Green 2009; Webb et al. 2019). Three key characteristics of the circadian clocks used to distinguish circadian-regulated phenotypes from other responses to environmental cues are (1) the rhythm has an approximate 24-hour period, (2) the rhythms can be reset by the environmental signals, and (3) the rhythmic period is consistent across a set of temperature ranges (McClung 2006). The simplified clock model comprises inputs, central oscillators, and outputs (Harmer 2009). When plants perceive changes in the inputs such as light and temperature, the core oscillators adjust their expression to regulate rhythmic outputs such as leaf movement, stomatal opening, hypocotyl elongation, and flowering time (Barak et al. 2000; Dowson-Day and Millar 1999; Engelmann, Simon, and Phen 1992; Somers et al. 1998). Arabidopsis clock proteins form complex transcription-translation feedback loops (Nakamichi 2011; Harmer 2009). Key proteins in these circadian loops are CIRCADIAN CLOCK ASSOCIATED 1 (CCA1), LATE ELONGATED HYPOCOTYL (LHY), and TIMING OF CAB EXPRESSION 1 (TOC1) (Alabadí et al. 2001; Strayer et al. 2000). The morning-expressed proteins CCA1 and LHY redundantly repress the expression of the evening gene, *TOC1* (Alabadí et al. 2001) and vice versa (Perales and Más 2007; Pruneda-Paz et al. 2009a). Another key component in the clock system is the evening complex (EC) which consists of LUX ARRHYTHMO (LUX), EARLY FLOWERING 3 (ELF3), and ELF4 (Nusinow et al. 2011a). This complex accumulates at dusk and represses the expression of *PSEUDO RESPONSE REGULATOR9* (*PRR9*), a CCA1/LHY repressor, creating a negative feedback loop with the morning genes (Dixon et al. 2011; Chow et al. 2012; Helfer et al. 2011; Herrero et al. 2012). The clock components directly or indirectly interact with each other, resulting in a complex network to control output genes in the circadian-related pathways such as CHLOROPHYLL A/B BINDING PROTEIN (CAB2) in photosynthesis (Kay 1993; Millar, Carré, et al. 1995), FLAVIN-BINDING, KELCH REPEAT, F-BOX PROTEIN 1 (FKF1) in flowering (Imaizumi et al. 2003), and PHYTOCHROME INTERACTING FACTOR4 (PIF4) in plant growth (Choi and Oh 2016).

ELF3, a component of the EC, is a multifunctional protein involving both light and temperature signaling (McWatters et al. 2000a; M. F. Covington et al. 2001a; Hicks, Albertson, and Wagner 2001; Jung et al. 2020). ELF3 regulates the gating of light inputs to the clock and light resetting of the clock (McWatters et al. 2000a; M. F. Covington et al. 2001a; Hicks, Albertson, and Wagner 2001). Through interaction with PHYTOCHROME B (PHYB), ELF3 modulates light input to the clock (Kolmos et al. 2011; Liu et al. 2001). ELF3 is required for temperature entrainment; the *elf3* mutant maintains rhythmic expression in thermocycles but does not exhibit a 24-hour circadian rhythm in either constant light or temperature (Thines and Harmon 2010). In addition to arrhythmia in free-running conditions, loss of ELF3 has several phenotypes, including the lack of a light gating response and altered photomorphogenesis (McWatters et al. 2000a; M. F. Covington et al. 2001a; Hicks, Albertson, and Wagner 2001; Zagotta et al. 1996). ELF3 also regulates periodic flowering by interacting with CONSTITUTIVE PHOTOMORPHOGENIC 1 (COP1) to control the stability of GIGANTEA (GI), which is a part of the flowering pathway (Yu et al. 2008). ELF3 affects hypocotyl elongation in response to both light and temperature. ELF3 is instrumental in gating hypocotyl elongation at dusk through interactions with the light-signaling proteins PHYB and COP1 and, as a component of the evening complex, by repressing *PIF4* (Nieto et al. 2015; Nusinow et al. 2011b; Yu et al. 2008; Liu et al. 2001). As PIF4 is a center of thermoresponsive growth, ELF3 is also involved in thermomorphogenesis by gating *PIF4* expression in response to temperature elevation (Box et al. 2015; Raschke et al. 2015). A prion-like domain (PrD) of ELF3 functions as a thermosensor (Jung et al. 2020). In Arabidopsis, a variation in the length of the polyQ region inside the PrD results in different thermal responsiveness, but removing the PolyQ region does not entirely remove thermal responsiveness, suggesting that the whole PrD is essential for temperature response (Jung et al. 2020). ELF3 with no PrD results in no thermal responsiveness, and the PrD controls the temperature-dependent conformation of ELF3 (Jung et al. 2020). These multiple roles of ELF3 in circadian regulation of light and temperature signaling pathways suggest that it is an integrator between the clock and environmental cues.

Understanding the cellular clock is complicated by intercellular communication. In plants, local (tissue-specific) clocks and cells communicate as a hierarchical network to synchronize circadian rhythms at the organismal level (Sorkin and Nusinow 2021; Takahashi et al. 2015). In humans, the suprachiasmatic nuclei (SCN) serve as a pacemaker to synchronize the clocks in peripheral organs via electrical, endocrine, and metabolic signaling pathways (Albrecht 2012). However, the structure of plant circadian rhythms is still under debate. One model is the hierarchical network, similar to mammals, where the shoot apical meristem is a dominant clock that transmits circadian information to roots (Takahashi et al. 2015). Root and shoot clocks behave differently in the presence of light but act similarly in darkness (Bordage et al. 2016). The EC involves the difference between shoot and root clocks in response to light quality as roots and shoots of *elf3*, *elf4*, and *lux* mutants show different rhythmicity and period (Nimmo et al. 2020). ELF4 is also essential for controlling the pace of the root clock in response to temperature (W. W. Chen et al. 2020). Another model is a decentralized model where the local clocks equally contribute to the rhythms in the whole plant (Endo 2016). The evidence supporting this idea is that root tips with high cell density exhibit strong local coupling between cells like shoot apical meristem (Gould et al. 2018; Takahashi et al. 2015). It is proposed that plants have multiple signaling hubs to synchronize the whole-plant oscillations (Sorkin and Nusinow 2021). As callus is a unique type of plant tissue composed of homogeneous cells, we were interested in determining the circadian rhythms in callus.

Arabidopsis cell suspension cultures have been successfully used to investigate the components and kinetics of the Arabidopsis circadian clock, supporting the idea that this system could be a powerful tool for understanding intracellular circadian regulation (W.-Y. Kim, Geng, and Somers 2003). Callus systems have been used to generate transgenic reporters in brassica, allowing a quick evaluation of many genotypes compared to transforming intact plants (Xu, Xie, and McClung 2010). Due to the ability of callus to be converted to an embryo-like structure and regenerated to a new type of cells (Jha 2005; Krikorian and Berquam 1969), callus is also used in micropropagation (El-Esawi 2016), and the generation of transgenic plants with desired traits (Y.-H. Kim et al. 2011; Rai et al. 2011; Wu et al. 2015). To induce callus from explants in vitro, two plant growth regulators, auxin and cytokinin, are used (Grafi and Barak 2015). However, callus systems have different gene expression profiles than other tissues (Tanurdzic et al. 2008; He et al. 2012; Du et al. 2019; Shim et al. 2020; K. Lee, Park, and Seo 2018). About 138 circadian-related genes, including *CCA1*, *LHY*, *TOC1*, and *ELF3,* were differentially expressed in Arabidopsis callus compared to intact leaves, suggesting that changes in clock gene expression were important for the establishment and maintenance of callus (K. Lee, Park, and Seo 2016).

Even though callus and cell suspension cultures are used in basic research and industrial application, reports vary on the similarity of the circadian rhythm between intact plants and callus and cell suspension systems (W.-Y. Kim, Geng, and Somers 2003; Nakamichi et al. 2003, 2004; Xu, Xie, and McClung 2010; Jiqing Sai and Johnson 1999). *Brassica napa* callus is considered to have functional clocks based on the observation that *CCA1* expression was entrained by both photocycles and thermocycles (Xu, Xie, and McClung 2010). However, in these calli, the period of CCA1 was consistently longer than the cotyledon movement period in intact seedlings. Tobacco callus exhibited the rhythmic oscillation of the *LiGHT-HARVESTING CHLOROPHYLL A/B-BINDING (LHCB)* gene in constant light, indicating that it retains a functional circadian clock (J. Sai and Johnson 1999). In Arabidopsis, the cell suspension T87 had robust circadian clocks as the expression of *CCA1* and *TOC1* was rhythmic under the constant light and dark similar to intact plants (Nakamichi et al. 2003, 2004). Although another line of Arabidopsis cell suspension also maintained rhythmic *CCA1* and *TOC1* mRNA levels under free-running conditions, the period was significantly more extended than in intact plants (W.-Y. Kim, Geng, and Somers 2003).

This study aimed to characterize the circadian oscillation in the Arabidopsis callus obtained from the cell suspension named AT2 (Tanurdzic et al. 2008)). Examining multiple transcriptional reporters, we found that the AT2 callus lacked circadian oscillations in constant light and had an altered period compared to intact seedlings. We found that the loss of oscillations in constant light is specific to the AT2 callus, but the lengthened period was also observed in freshly derived callus. We then determined the possible factors contributing to altered oscillations in this callus and freshly generated callus. Finally, we determined the effect of ELF3 overexpression on circadian oscillations in the AT2 callus.

## 2. Materials and Methods

### 2.1 Plant growth conditions

Arabidopsis AT2 cell suspension culture was provided by Linda Hanley-Bowdoin, North Carolina State University, Raleigh, NC, USA. This cell line was established from *Arabidopsis thaliana* ecotype Columbia-0 (Col-0) (Tanurdzic et al. 2008). Cells were maintained as described in (Tanurdzic et al. 2008) with small modifications. Cells were grown in Gamborg’s B5 (GB5) media (Gamborg, Miller, and Ojima 1968) supplemented with 2.5 mM MES, 0.5 μg/ml 2,4-D, and 3% (w/v) sucrose (pH 5.8) (Tanurdzic et al. 2008). Cells were shaken at 160 rpm under 12/12 h light/dark cycles (LD) and subcultured every week by transferring 5 ml cell suspension to 50 ml fresh media.

*CCA1::LUC (Pruneda-Paz et al. 2009b)*, *LHY::LUC (Baudry et al. 2010)*, *TOC1::LUC (Alabadí et al. 2001)*, and *FKF1::LUC* seeds were sterilized and plated on ½ MS media (Murashige and Skoog 1962) with 0.4% (w/v) agar. Seeds were stratified at 4°C for three days and then moved to a growth chamber set to 23°C and LD. Seedlings (1-2 weeks old) were used in the time-course bioluminescence assay.

### 2.2 *Agrobacterium* transformation and initiation of AT2 transformed callus

The seven-day-post-subculture AT2 cells were pre-cultured in transformation media (GB5 media supplemented with 2.5 mM MES, 2.7 μM NAA, 0.23 μM kinetin, and 3% (w/v) sucrose) for 2 days. *Agrobacterium tumefaciens* strain GV3101 containing *CCA1::LUC*, *LHY::LUC*, *TOC1::LUC*, and *FKF1::LUC* were co-cultured with AT2 cells supplemented with 100 μM acetosyringone for two days. Transfected cells were then washed and grown in transformation media containing 250 μg/ml timentin with shaking for three days. Transformed cells were plated on selective media (GB5 media supplemented with 2.5 mM MES, 2.7 μM NAA, 0.23 μM kinetin, 250 μg/mL timentin, 3% (w/v) sucrose, and 0.8% (w/v) agar) for two weeks. The 50 μg/ml gentamicin was used to select *CCA1::LUC*, *LHY::LUC*, *TOC1::LUC*, and *FKF1::LUC*. Selected calli were transferred to fresh selective agar plates every two weeks to maintain their growth. Transformed calli used in the time-course bioluminescence assay were grown on agar plates for 7-12 days after the previous subculture.

### 2.3 Generating the overexpression of ELF3 in AT2 transformed callus

Gateway cloning technology was used to generate the *35s*::*ELF3* construct. The *ELF3* coding sequence (CDS) in the pENTR vector (Pruneda-Paz et al. 2014) was cloned into the 35s-promoter pB7WG2 vector (Karimi, Inzé, and Depicker 2002) via LR reaction (Gateway LR Clonase II Enzyme mix, Invitrogen, USA). *A. tumefaciens* strain GV3101 carrying *35s::ELF3* vector was used to infect *CCA1::LUC*, *LHY::LUC*, and *TOC1::LUC* AT2 calli as described above. Transformed calli containing both *35s*::*ELF3* and clock reporters were selected on media supplemented with 50 μg/mL gentamicin and 10 μg/mL glufosinate-ammonium for two weeks. Selected calli were subcultured to fresh selection media every two weeks.

### 2.4 Callus induction from Arabidopsis tissues and seedlings

*CCA1::LUC*, *LHY::LUC*, and *FKF1::LUC* seeds were sterilized in 50% (v/v) bleach and 0.02% (v/v) Triton X-100 and placed on callus induction media. We used two recipes of callus induction media reported in (Barkla, Vera-Estrella, and Pantoja 2014) and (Sello et al. 2017). The Barkla media was MS media supplemented with 1 µg/ml 2,4-D, 0.05 µg/ml BA, 3% (w/v) sucrose, and 0.8% (w/v) agar (pH 5.7) (Barkla, Vera-Estrella, and Pantoja 2014). The Sello media was MS media supplemented with 0.5 µg/ml 2,4-D, 0.25 µg/ml BA, 3% (w/v) sucrose, and 0.8% plant agar (pH 5.5) (Sello et al. 2017). Seeds were stratified at 4°C for three days, and the plates were then moved to a growth chamber with 23°C and LD. Callus was generated from seedlings within three weeks. Calli were subcultured to fresh media every two weeks to maintain their growth.

To generate calli from different plant tissues, *CCA1::LUC* seeds were sterilized as described above and placed on ½ MS media with 0.4% (w/v) agar. Seeds were stratified at 4°C for three days and then moved to a growth chamber with the condition described above. Hypocotyls, first leaves, and roots were isolated from seven-day-old seedlings and placed on the Barkla callus induction media (Barkla, Vera-Estrella, and Pantoja 2014). Calli induced from those tissues were subcultured to fresh media every two weeks.

### 2.5 Luciferin spraying and time-course bioluminescence assay

Calli and seedlings were sprayed with 250 μg/ml D-luciferin (GoldBio, USA) in 0.01% (v/v) Triton X-100. The plates were placed in the growth chamber installed with a CCD camera (Eagle V 4240 Scientific CCD camera, Raptor Photonics). The camera was controlled by Micro-Manager software (Edelstein et al. 2014), and images were captured with a 20-minute exposure time every two hours to detect bioluminescence. For the LD condition, the light was on at 8 am (circadian time 0; ZT0) and off at 8 pm (ZT12). To transition from LD to the free-running conditions, the lights were kept on starting at 8 pm (ZT12) for continuous light (LL), and the lights were kept off starting at 8 am (ZT0) for continuous darkness (DD). In the temperature cycles (HC), daytime temperatures were kept at 23°C for 12 h and nighttime temperatures were 12°C.

The luminescence signal of luciferase from each callus or seedling was measured as mean gray values (total intensity in the selected area divided by the number of pixels) using Image J software (Schneider, Rasband, and Eliceiri 2012). The mean gray value was subtracted by the mean gray value of the background to obtain the normalized intensity. To calculate a relative normalized intensity, the normalized intensity at individual time points was divided by a maximum normalized intensity in that time series. Relative normalized intensity was then linearly detrended on BioDare2 (biodare2.ed.ac.uk) (Zielinski et al. 2014). Rhythmicity test on BioDare2 was used to select only rhythmic data for making time-series plots (Hutchison et al. 2015). Periods and amplitudes were estimated by a Fourier transform–non-linear least squares (FFT-NLLS) method on BioDare2 (Zielinski et al. 2014). Plots were made in R version 4.0.5.

Statistical analysis was performed in R version 4.0.5 to compare periods and amplitudes of bioluminescence signals. Two-sample Wilcoxon rank-sum test was used to compare means between calli and seedlings in the same conditions or between calli on different media. One-way ANOVA was used to compare the means between calli derived from different tissues.

### 2.6 RNA-Seq data processing

We obtained RNA-Seq data of AT2 cell suspension from Dr. Linda Hanley-Bowdoin (PRJNA412215 and PRJNA412233 on NCBI SRA) and RNA-Seq data of Col-0 seedlings from (Grinevich et al. 2019) (PRJNA488799 on NCBI SRA) to compare the read counts of clock genes between the AT2 cell suspension and seedlings. The cell suspension was maintained as described in (T.-J. Lee et al. 2010). The AT2 cell suspension was grown in LL and DD for seven days and the cells were then harvested with three biological replicates. However, the time of day the cells were harvested was not reported. The Col-0 seedlings used for comparison were grown in LL and harvested at dawn and dusk as described in (Grinevich et al. 2019). FastQC (version 0.11.8) (Andrews 2010) (“Babraham Bioinformatics - FastQC A Quality Control Tool for High Throughput Sequence Data” n.d.) was used to quality check reads before and after adapter trimming by BBDuk (in BBMap version 38.34; https://sourceforge.net/projects/bbmap/) (Bushnell 2014; “BBMap Guide” 2016). Reads were aligned to Arabidopsis reference genome TAIR10 (Berardini et al. 2015) using Hisat2 (version 2.2.0) (D. Kim et al. 2019). Read counts were obtained by featureCounts from the Rsubread package (version 2.0.1) (Liao, Smyth, and Shi 2014). Low read counts were filtered out by the command *filterByExpr()* from EdgeR (M. D. Robinson, McCarthy, and Smyth 2010; McCarthy, Chen, and Smyth 2012). The read counts were normalized by the median of ratio method in DESeq2 (version 1.26.0) to obtain normalized gene expression (Love, Huber, and Anders 2014). The heat maps of normalized read counts and percent rank of gene expression were generated using the R package “pheatmap” in R version 3.6.3.

### 2.7 Identification of SNPs in the AT2 cell suspension

Whole-genome reads were aggregated together using the following studies that use *A. thaliana* suspension culture (PRJNA412215 and PRJNA412233 on NCBI SRA). STAR (version 2.5.3a) was used to align reads. STAR’s 2-pass mapping was used with default parameters (Dobin et al. 2013). Picard (version 2.10.2; https://github.com/broadinstitute/picard) was used to order bam files and mark duplicate reads. Finally, GATK’s HaplotypeCaller (version 3.7; https://software.broadinstitute.org/gatk/) was used to call variants. Variants were filtered with FS > 30 and QD > 2. Sequence alignment was visualized on IGV version 2.8.13 (J. T. Robinson et al. 2011).

### 2.8 RNA extraction and quantitative real-time PCR (qRT-PCR)

To compare the expression of clock genes between the AT2 cell suspension and intact plants, seven-day-old AT2 cell suspension and two-week-old Col-0 seedlings were harvested at ZT8 and ZT12 and flash-frozen in liquid nitrogen before storing at -80°C. To compare the expression of clock genes between LD and LL in the AT2 callus, seven-day-old AT2 callus were harvested at ZT0 and ZT12. Total RNAs were extracted from 100 mg of plant tissues by RNeasy Plant RNA Mini Kit (Qiagen, USA). One nanogram of ArrayControl™ RNA Spikes (Invitrogen, USA) was added during tissue homogenization. DNase I treatment (Roche, USA) was performed on columns to remove DNA. The total RNA was measured by the NanoDrop™ Lite Spectrophotometer (Thermo Fisher, USA). For comparing gene expression between AT2 cell suspension and Col-0, 500 ng of total RNA were used in cDNA synthesis with iScript™ Advanced cDNA Synthesis Kit for RT-qPCR (Bio-Rad, USA) while 700 ng of total RNA from the AT2 callus in LD and LL were used as an input. cDNAs were then 1:4 diluted in nuclease-free water prior to the qPCR. The qPCR reaction contained 5 µl of iTaq™ Universal SYBR® Green Supermix (Bio-Rad, USA), 1 µl of 5 µM forward primer, 1 µl of 5 µl reverse primer, 2 µl diluted cDNA, and 1 µl of nuclease-free water. The qPCR was performed on the CFX384 Touch Real-Time PCR Detection System (Bio-Rad, USA) under the following conditions: initial denaturation at 95°C for 30 seconds and the 35 cycles of denaturation (95°C for 5 seconds) and annealing (60°C for 30 seconds). Melt curve analysis (65 to 95°C with a 0.5°C increment) was done at the end of qPCR cycles. Only samples with a single discrete peak in the melt curve analysis were kept for the gene expression analysis. Four technical replicates were performed for each cDNA sample in each target gene. Relative normalized expression (ΔΔCq) was determined by CFX Maestro 1.1 software (Bio-rad, USA). External RNA spikes and three housekeeping genes *ISOPENTENYL DIPHOSPHATE ISOMERASE 2* (*IPP2*), *ACTIN 2* (*ACT2*), and *TUBULIN 2/3* (*TUB2/3*) were used as reference genes to compare gene expression between the AT2 cell suspension and intact plants. Only external RNA spikes and *IPP2* were used as references to compare gene expression between LD and LL in the AT2 callus. Primers used in the study were listed in Supplementary Table 1. The two-sample t-test was used to compare the means between two time points. Statistical analysis and making plots were performed in R version 4.0.5.

## 3. Result

### 3.1 AT2 callus showed a rhythmic expression in LD but not in the free-running condition

We aimed to observe whether the Arabidopsis AT2 callus has a functional circadian clock by generating the calli carrying *CCA1::LUC*, *LHY::LUC*, *TOC1::LUC*, and *FKF1::LUC* via *Agrobacterium*-mediated transformation. Gentamicin was used to select transformants. To ensure that the changes we observed were not due to the selective antibiotic, we compared the expression of *CCA1::LUC*, *LHY::LUC*, *TOC1::LUC*, and *FKF1::LUC* in the AT2 callus cells grown on selective media and regular media under LD (Supplementary Figure 1). There was no significant difference in waveforms, and the expression of clock genes in calli on both media had an approximately 24-h period (Supplementary Figure 1 and Table 1). This indicated that gentamicin did not affect the expression in LD of the circadian clock genes. Therefore, we decided to maintain the AT2 callus on gentamicin in further experiments.

**Table 1.** Periods of clock genes in the AT2 calli grown with and without gentamicin compared to periods of clock genes in seedlings under LD for 60 hours. Period lengths, represented as mean ± SD, are based on luciferase reporters for each of the listed clock-associated genes. n refers to the number of individual callus groups or seedlings. Callus with no gentamicin: Two-sample Wilcoxon rank-sum test was used to compare the means between the callus with and without gentamicin; all reporters showed no significant (ns) differences between growth with and without gentamicin, P > 0.05. Seedlings: Two-sample Wilcoxon rank-sum test was used to compare the means between the AT2 callus with gentamicin and seedlings. The significance levels are in the ‘seedlings’ column; ns = not significant (P> 0.05), * = significant at P ≤ 0.05, ** = significant at P ≤ 0.01, *** = significant at P ≤ 0.001, and **** = significant at P ≤ 0.0001.

| Gene | Callus with gentamicin | Callus with no gentamicin | Seedlings |
| --- | --- | --- | --- |
| *CCA1::LUC* | 24.09 ± 0.70 (n=34) | 24.09 ± 0.41 (n=36) ns | 23.68 ± 0.28 (n=32) **** |
| *LHY::LUC* | 24.05 ± 0.32 (n=18) | 24.23 ± 0.25 (n=35) ns | 24.02 ± 0.24 (n=28) ns |
| *TOC1::LUC* | 24.69 ± 0.93 (n=17) | 24.72 ± 0.99 (n=36) ns | 24.31 ± 0.33 (n=17) * |
| *FKF1::LUC* | 24.66 ± 0.13 (n=18) | 24.63 ± 0.34 (n=36) ns | 24.39 ± 0.21 (n=12) *** |

The generation of callus and cell suspension culture requires genetically and epigenetically reprogramming (Tanurdzic et al. 2008; He et al. 2012; Ikeuchi, Sugimoto, and Iwase 2013; Du et al. 2019; Shim et al. 2020). Plant growth regulators are the key to switch cell fate in tissue culture systems (Grafi and Barak 2015). Other factors such as light (Batista et al. 2018), humidity (C. Chen 2004), and osmotic potential (de Paiva Neto and Otoni 2003) in tissue culture could induce stress in cultured tissues, leading to changes in genetics and epigenetics (Bednarek and Orłowska 2020). These unnatural environments could also affect the rhythms in the AT2 callus compared to intact plants. In addition, if coordination across tissues is required for coordinated expression and proper timing of gene expression in plants, the callus with limited cellular organization, may have disruptions in their rhythmic expression of circadian and clock-controlled genes. Therefore, we examined the expression of *CCA1::LUC*, *LHY::LUC*,

*TOC1::LUC*, and *FKF1::LUC* in the AT2 callus cells and intact seedlings under LD (Figure 1 and Supplementary Figure 2). The expression waveforms of *CCA1::LUC* and *LHY::LUC* in the AT2 callus cells under the LD were similar to those in intact plants (Figure 1A, 1B, Supplementary Figure 2A, 2B). However, The peaks of *TOC1::LUC* and *FKF1::LUC* in AT2 callus cells were broader than those in seedlings (Figure 1D, 1E, Supplementary Figure 2D, and 2E). The periods of *CCA1::LUC*, *TOC1::LUC*, and *FKF1::LUC* in AT2 callus cells under LD were significantly longer than those in seedlings, but there was no significant difference in the *LHY*::*LUC* period between the AT2 callus and seedlings (Table 1). Even though several clock genes in the AT2 callus had longer periods than those in seedlings, their periods were still in the 24-h range (Table 1). This suggested that, while under entraining conditions of light/dark cycles, the overall cascade and timing of clock gene expression in the AT2 callus was similar to that in seedlings.

**Figure 1.**
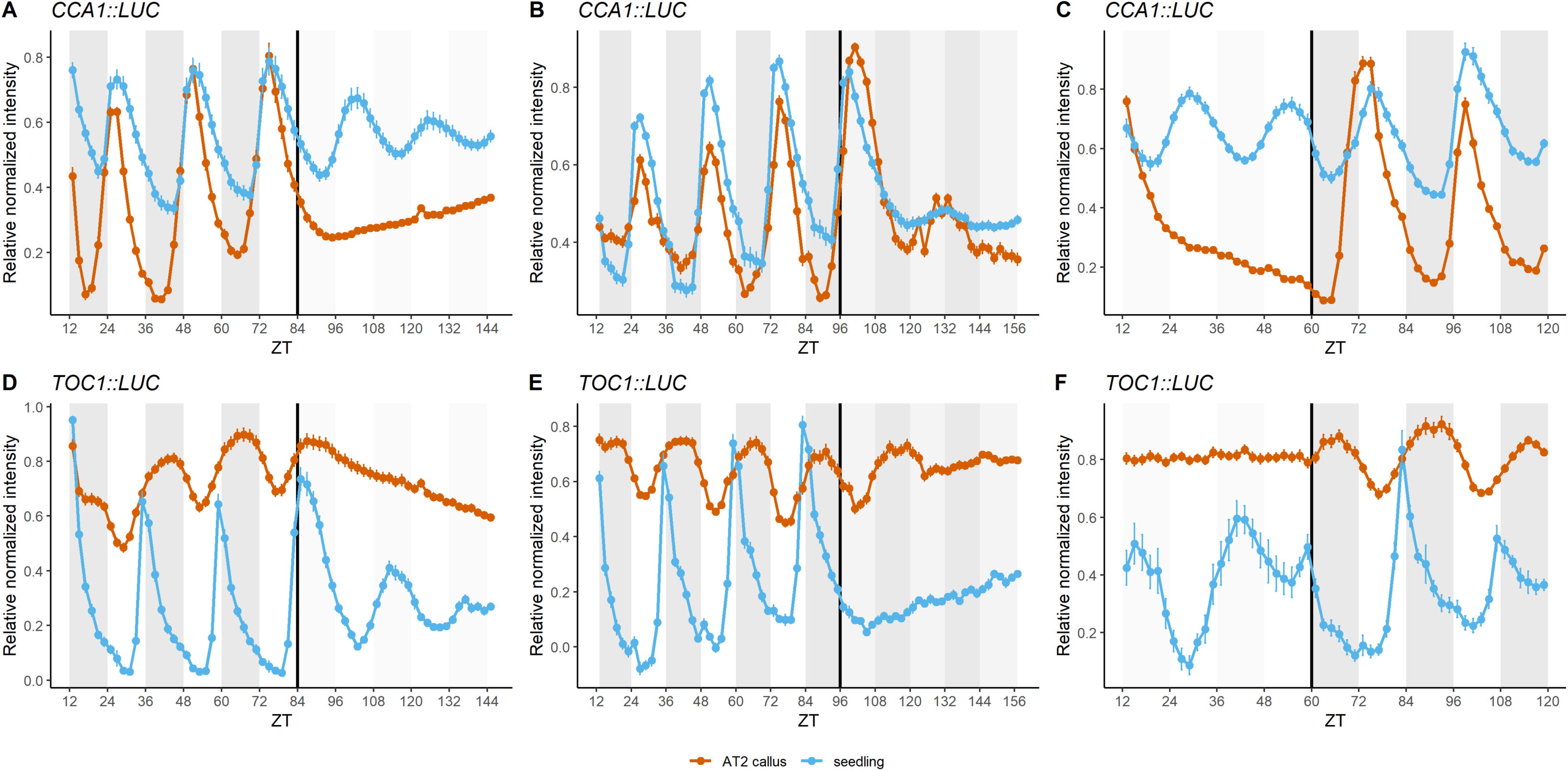
The AT2 callus had an altered circadian oscillation. Bioluminescence assay of *CCA1::LUC* (A-C) and *TOC1::LUC* (D-F) in the AT2 callus and seedlings under LD for 72 hours and subsequent LL for 60 hours (A and D), LD for 84 hours and subsequent DD for 60 hours (B and E), and LL for 48 hours and subsequent LD for 58 hours (C and F). Data are mean ± SEM. n = 14-36 individual callus groups and n = 9-32 seedlings, see Tables 2, 3 and Supplemental Table 2 for specific n for each reporter.

**Table 2.** Periods and amplitudes of clock genes, represented as mean ± SD, are shown for the AT2 callus under LD and subsequent DD for 60 hours. n refers to a number of individual callus groups. A paired-sample Wilcoxon test was used to compare the means between LD and DD; ns = not significant, * = significant at P ≤ 0.05, ** = significant at P ≤ 0.01, *** = significant at P ≤ 0.001, and **** significant at P ≤ 0.0001.

| Gene | Period (hours). | Amplitude |
| --- | --- | --- |

**Table 2.** Periods and amplitudes of clock genes, represented as mean ± SD, are shown for the AT2 callus under LD and subsequent DD for 60 hours.
|  | LD | DD | LD | DD |
| --- | --- | --- | --- | --- |
| *CCA1::LUC* | 24.16 ± 0.58 (n=13) | 27.30 ± 1.17 ** | 0.2592 ± 0.0284 (n=13) | 0.1208 ± 0.0293 ** |
| *LHY::LUC* | 23.76 ± 0.32 (n=28) | 28.99 ± 1.39 **** | 0.2418 ± 0.0370 (n=28) | 0.0757 ± 0.0235 **** |
| *TOC1::LUC* | 24.33 ± 0.54 (n=11) | 26.96 ± 1.23 ** | 0.1882 ± 0.0260 (n=11) | 0.0655 ± 0.0113 ** |
| *FKF1::LUC* | 24.43 ± 0.13 (n=27) | 29.05 ± 0.59 **** | 0.2641 ± 0.0241 (n=27) | 0.1526 ± 0.0296 **** |

**Table 3.** Periods of the clock reporters are represented as mean ± SD in AT2 callus cells and seedlings in LD for 58 hours after growth under LL for 48 hours. n refers to a number of individual callus groups or seedlings. Two-sample Wilcoxon rank-sum test was used to compare the means between the calli and seedlings; ns = not significant (P > 0.05), * = significant at P ≤ 0.05, ** = significant at P ≤ 0.01, *** = significant at P ≤ 0.001, and **** = significant at P ≤ 0.0001.

| Gene | AT2 callus | Seedlings |
| --- | --- | --- |
| *CCA1::LUC* | 25.66 ± 0.30 (n=36) | 24.82 ± 0.33 (n=9) **** |
| *LHY::LUC* | 25.65 ± 0.54 (n=18) | 26.13 ± 1.43 (n=6) ns |
| *TOC1::LUC* | 25.97 ± 1.24 (n=18) | 25.58 ± 0.66 (n=9) ns |
| *FKF1::LUC* | 28.46 ± 1.47 (n=18) | 25.06 ± 0.80 (n=8) *** |

Circadian-controlled rhythms are sustained under free-running conditions (Somers et al. 1998; Dowson-Day and Millar 1999; Engelmann, Simon, and Phen 1992; Harmer 2009). To determine if the differences in the waveforms we observed in LD conditions affected the rhythmic expression in free-running conditions, we observed the bioluminescence of *CCA1::LUC*, *LHY::LUC*, *TOC1::LUC*, and *FKF1::LUC* in AT2 callus cells under continuous light (LL) and continuous dark (DD) at constant temperatures (Figure 1 and Supplementary Figure 2). In LL, there was no rhythmic oscillation in the AT2 callus in contrast to seedlings where the expression of *CCA1::LUC*, *LHY::LUC*, *TOC1::LUC*, and *FKF1::LUC* persisted albeit with significantly decreased amplitudes and longer periods (Figure 1A, 1D, Supplementary Figure 2A, 2D, and Table 2). The qRT-PCR results from the AT2 callus grown in LD and LL for seven days confirmed that *CCA1*, *LHY*, *TOC1*, *ELF4*, and *GI* significantly lost their rhythmicity under LL compared to LD (Supplementary Figure 3A, 3B, 3C, 3E, and 3G). In LD, these transcripts showed significant differences in expression between ZT0 and ZT12, but the differences in expression were not significant in LL (Supplementary Figure 3A, 3B, 3C, 3E, and 3G). Surprisingly, unlike the expected expression pattern in seedlings, *ELF3* showed no difference in expression between dawn and dusk in both LD and LL while *LUX* was differentially expressed only in LL (Supplementary Figure 3D and 3F). In LD, *LUX* tended to be up-regulated in the evening, but the difference was not significant (Supplementary Figure 3F). In summary, the rhythmic expression of the core clock genes in the AT2 callus was lost in LL.

We also observed the expression of clock genes in DD which is another type of free-running condition. In seedlings, the clock genes rapidly lost their rhythmicity under DD (Figure 1B, 1E, Supplementary Figure 2B, and 2E). The first expression peak of *CCA1::LUC*, *LHY::LUC*, and *FKF1::LUC* in DD occurred due to the preceding LD (Millar, Straume, et al. 1995). In seedlings, there was no second peak of *CCA1::LUC*, *LHY::LUC*, and *FKF1::LUC* after 24 hours in DD while the *TOC1::LUC* expression diminished right after the transition to DD. The circadian oscillation usually dampens faster in DD than in LL possibly due to reduced phototransduction to input the clocks in darkness (Millar, Straume, et al. 1995; C. H. Johnson et al. 1995). However, in the AT2 callus, the clock gene expression persisted under DD with a longer period and smaller amplitude than in LD conditions (Figure 1B, 1E, Supplementary Figure 2B, and 2E). Although the rhythmic expression in the AT2 callus was more pronounced in DD than LL and had a stronger rhythmic expression than seedlings in DD, the periods in the AT2 callus under DD were longer than the LL seedling periods (Table 2 and Supplemental Table 2). Therefore, although the AT2 callus maintained rhythmic expression in DD, this is a weak rhythm compared to the LL expression in seedlings. This indicates that the AT2 callus has a circadian oscillator that can function, albeit poorly, in free-running conditions, but the loss of rhythms in LL conditions suggests a disruption in the AT2 circadian oscillator compared to intact seedlings.

Since the AT2 callus displayed a rhythmic expression of core clock genes under LD and DD but not in the LL condition, it is possible that the light acts as a signal in LL to drive the expression. To examine this, we measured the bioluminescence of *CCA1::LUC*, *LHY::LUC*, *TOC1::LUC*, and *FKF1::LUC* transferred from LL to LD in the dark period (Figure 1C, 1F, Supplementary Figure 2C, and 2F). Seedlings were able to maintain the rhythmic expression in both LL and the subsequent LD. As observed above, there was no rhythmic expression in the AT2 callus under LL. However, the AT2 callus resumed rhythmic expression of these clock genes in the first dark period and immediately returned to the rhythmic waveforms observed in LD. For example, *CCA1::LUC* and *LHY::LUC* show a constant decrease in LL, but once in the dark, the expression levels gradually increase and peak in the light period (Figure 1C and Supplementary Figure 2C). However, the expression of evening genes *TOC1::LUC* and *FKF1::LUC* started rising right after the dark period (Figure 1F and Supplementary Figure 2F). The periods of all reporters were longer in the two days of LD after LL in both seedlings and calli compared to prior analysis of plants grown continuously in LD (Tables 1, 3, and Supplementary Table 2). The periods of *CCA1::LUC* and *FKF1::LUC* in the calli under the subsequent LD were significantly longer than those in seedlings while there was no significant difference in periods of *LHY::LUC* and *TOC1::LUC* between the calli and seedlings (Table 3). The recovery of the expected expression pattern in the dark suggests that light is disrupting the expression of clock promoters in the AT2 callus.

We found that the AT2 callus had altered circadian oscillation in LL. However, several publications show that the circadian clocks run normally in callus (W.-Y. Kim, Geng, and Somers 2003; Nakamichi et al. 2003, 2004; Xu, Xie, and McClung 2010; Jiqing Sai and Johnson 1999). We wondered whether the altered circadian rhythms were specific to the AT2 callus. We induced calli from *CCA1::LUC, LHY::LUC,* and *FKF::LUC* seedlings on two recipes of callus induction media, one from (Barkla, Vera-Estrella, and Pantoja 2014) and another from (Sello et al. 2017). Both recipes used the same plant growth regulators (2,4-D as auxin and BA as cytokinin), but at different concentrations. We measured the bioluminescence under LD and found that callus cells induced by both media exhibited similar rhythmicity within a 24-hour period (Figure 2 and Table 4). The period of *CCA1::LUC* in callus cells on two media under LD was not significantly different, but the periods of *LHY::LUC* and *FKF1::LUC* on the Sello media were significantly longer than on Barkla media (Table 4). We observed the expression of clock genes under LL to determine whether the oscillations persisted in these calli (Table 4). Calli derived from both recipes exhibited sustained oscillations in LL (Figure 2 and Table 4), suggesting the plant growth regulators used for callus induction did not result in a loss of rhythmicity of these clock-associated promoters. The periods of *LHY::LUC* and *FKF1::LUC* in callus cells under LL were significantly longer than those under LD in both media (Table 4). The *CCA1::LUC* period was longer in LL than LD only in the Sello media (Table 4). The Barkla media contained 1 µg/ml 2,4-D and 0.05 µg/ml BA, so the auxin-to-cytokinin ratio was 1:20 (Barkla, Vera-Estrella, and Pantoja 2014). The Sello media contained lower auxin but higher cytokinin (0.5 µg/ml 2,4-D and 0.25 µg/ml BA) than the Barkla media, and the ratio of auxin to cytokinin was 1:2 (Sello et al. 2017). Compared to the clock gene expression in the AT2 callus, and consistent with prior publications, the results indicated that the loss of circadian rhythms under LL in the AT2 callus is not inherent to the hormones used to generate the callus tissue.

**Figure 2.**
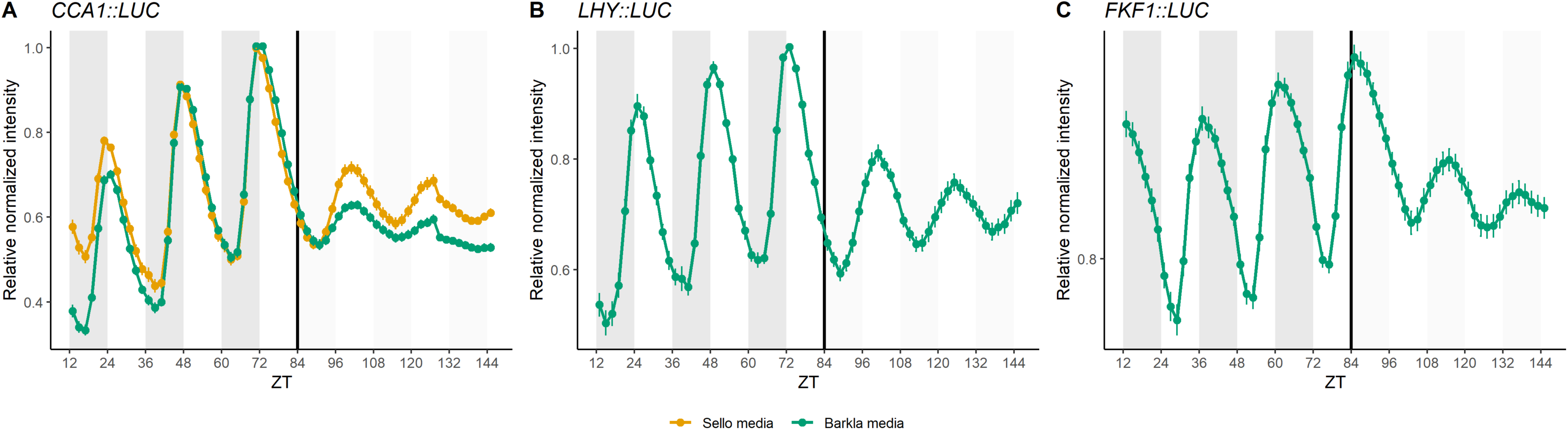
Callus derived from seedlings had normal circadian oscillation. Bioluminescence assay of (A) *CCA1*::*LUC*, (B) *LHY*::*LUC*, and (C) *FKF1*::*LUC* in calli derived from seedlings on the Sello (yellow) and the Barkla media (green) under LD for 72 hours and subsequent LL for 60 hours (n = 12-24 individual callus groups). Data are mean ± SEM.

**Table 4.** Period lengths of clock-associated reporters, represented as mean ± SD, are presented for calli derived from seedlings and grown on Sello and Barkla media under LD and LL for 60 hours. n refers to a number of individual callus groups. Two-sample Wilcoxon test was used to compare the means in LD between the Sello and Barkla media. The significance levels are shown in the ‘Barkla media-LD’ column; ns = not significant (P > 0.05), † = significant at P ≤ 0.01, and ‡ = significant at P ≤ 0.0001. Paired sample Wilcoxon test was used to compare the means between LD and LL. The significance levels are shown in the ‘LL’ columns; ns = not significant (P > 0.05), * = significant at P ≤ 0.05, ** = significant at P ≤ 0.01,*** = significant at P ≤ 0.001, and **** = significant at P ≤ 0.0001.

| Gene | Sello media |  | Barkla media |  |
| --- | --- | --- | --- | --- |
|  | LD | LL | LD | LL |
| *CCA1::LUC* | 24.26 ± 0.08 (n=12) | 24.91 ± 0.18 ** | 24.26 ± 0.13 (n=18) ns | 24.06 ± 0.73 ns |
| *LHY::LUC* | 24.30 ± 0.17 (n=18) | 24.76 ± 0.76 * | 24.09 ± 0.18 (n=16) † | 24.78 ± 0.59 *** |
| *FKF1::LUC* | 24.38 ± 0.21 (n = 18) | 30.86 ± 1.95 *** | 24.11 ± 0.18 (n = 24) ‡ | 25.62 ± 0.43 **** |

### 3.2 Temperature cycles can recover the circadian oscillations of clock-associated reporters in the AT2 callus under constant light conditions

Light and temperature are two primary signals that plants use to integrate their internal clock to environmental changes (Devlin and Kay 2001; McClung, Salomé, and Michael 2002). As constant light eliminated the rhythmic waveforms of clock genes in the AT2 callus, we wanted to determine if temperature cycles could recover rhythmic expression in LL. We measured the bioluminescence of *CCA1::LUC* and *FKF1::LUC* in the AT2 callus, seedlings, and callus derived from seedlings under constant 23°C (HH) and 23/12°C temperature cycles (HC) in LL (Figure 3A, 3B, 3D, and 3E). In seedlings and callus derived from seedlings, temperature cycles increased the amplitude of *CCA1::LUC*. In the AT2 callus, temperature cycles were able to recover the rhythmic expression of *CCA1::LUC* and *FKF1::LUC*. Additionally, in the AT2 callus, a small increase in *CCA1::LUC* level was detected when the temperature was shifted from warm to cool temperatures (23°C to 12°C) (Figure 3A). This small peak was not detectable in seedlings or callus derived from seedlings (Figure 3A and 3B). In contrast, *FKF1::LUC* expression was induced at the transition from cool to warm temperature (12°C to 23°C) in the AT2 callus, seedlings, and callus induced from seedlings (Figure 2D and 2E). These results indicate that entrainment by temperature cycles can recover the rhythmic expression of these core clock genes, although there are slight differences in their expression waveforms in the presence of these thermocycles.

**Figure 3.**
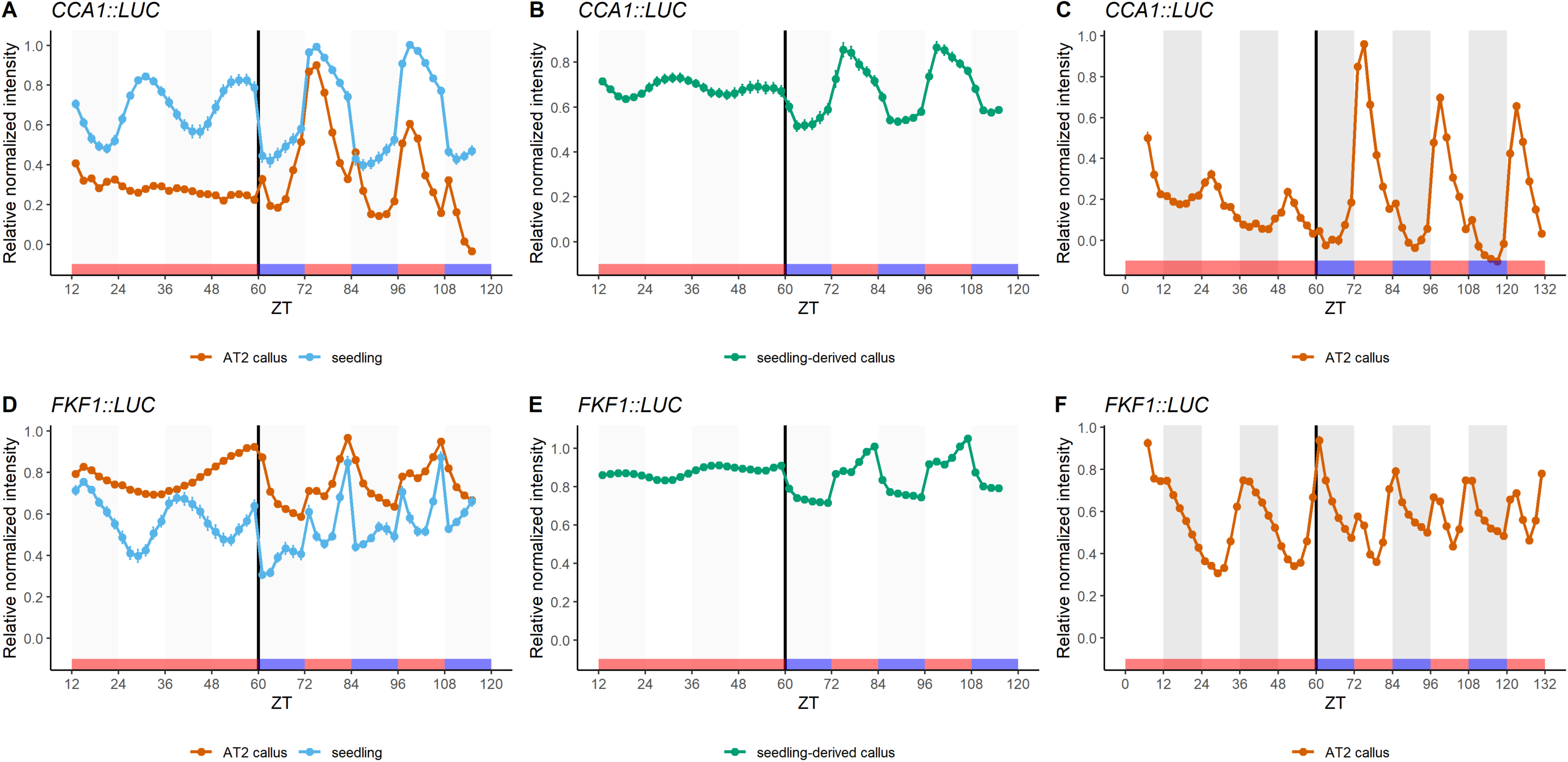
The AT2 callus showed a peak response to temperature changes. Bioluminescence assay of *CCA1::LUC* (A and B) and *FKF1::LUC* (D and F) in the AT2 callus (A and D) vs seedlings and calli derived from seedlings (B and E) on the Barkla media under LLHH for 48 hours and LLHC for 54 hours. Seedlings were entrained in LDHH for 14 days and then released to constant daytime temperature. Calli were entrained for 7 days (AT2 callus) or 9 days (callus derived from seedlings). Data are mean ± SEM. n = 26-36 individual AT2 callus groups, n= 13-30 seedlings, and n = 12-19 individual callus groups on the Barkla media. Bioluminescence assay of *CCA1*::*LUC* (C) and *FKF1*::*LUC* (F) in the AT2 callus under LDHH for 54 hours and LDHC for 70 hours. Data are mean ± SEM. n = 36 individual callus groups.

We next tested whether the temperature-responsive minor peaks observed in the AT2 callus persisted in the presence of both cycling light and temperatures by observing the expression of *CCA1::LUC* and *FKF1::LUC* under light and temperature cycles (LDHC) (Figure 3G and 3H). Both clock genes still showed two peaks, one ‘time of day’ peak, with a phase consistent with the single peak in LD conditions and a second, temperature-responsive peak in LDHC (Figure 3G and 3H). As the temperature-responsive peak appeared in both LD and LL, this suggested that this peak was specific to temperature changes and was not influenced by the constant light conditions. Overall, the expression of the clock genes examined in the AT2 callus showed that the rhythmic waveform not detectable in constant light could be recovered by the addition of temperature cycles.

### 3.3 Growth media and explant source did not contribute to the altered circadian rhythms in the AT2 callus

The AT2 callus was grown on maintenance media which contains different auxin and cytokinin (NAA and kinetin) compared to the media used to induce callus from seedlings (2,4-D and BA). As auxin and cytokinin affect the rhythmicity of clock genes in intact plants (Michael F. Covington and Harmer 2007; Zheng et al. 2006), We tested whether plant growth regulators used in callus induction media could recover the circadian oscillation in AT2 callus. We transferred the AT2 callus containing *CCA1::LUC* to the Barkla media and cultured them for a month. *CCA1::LUC* oscillation in the AT2 callus on both media was similar (Figure 4A). Even though the AT2 callus on the Barkla media showed a significant shortening period in LD compared to that in the AT2 callus on its original media, the periods were still in the 24-hour range (Table 5). This suggested that the Barkla media did not affect the *CCA1::LUC* expression in the AT2 callus in LD. In the constant light, the *CCA1::LUC* oscillation was not detectable in the AT2 calli on both media, indicating that the culture media was not the cause of the disrupted LL circadian oscillations in the AT2 callus.

**Figure 4.**
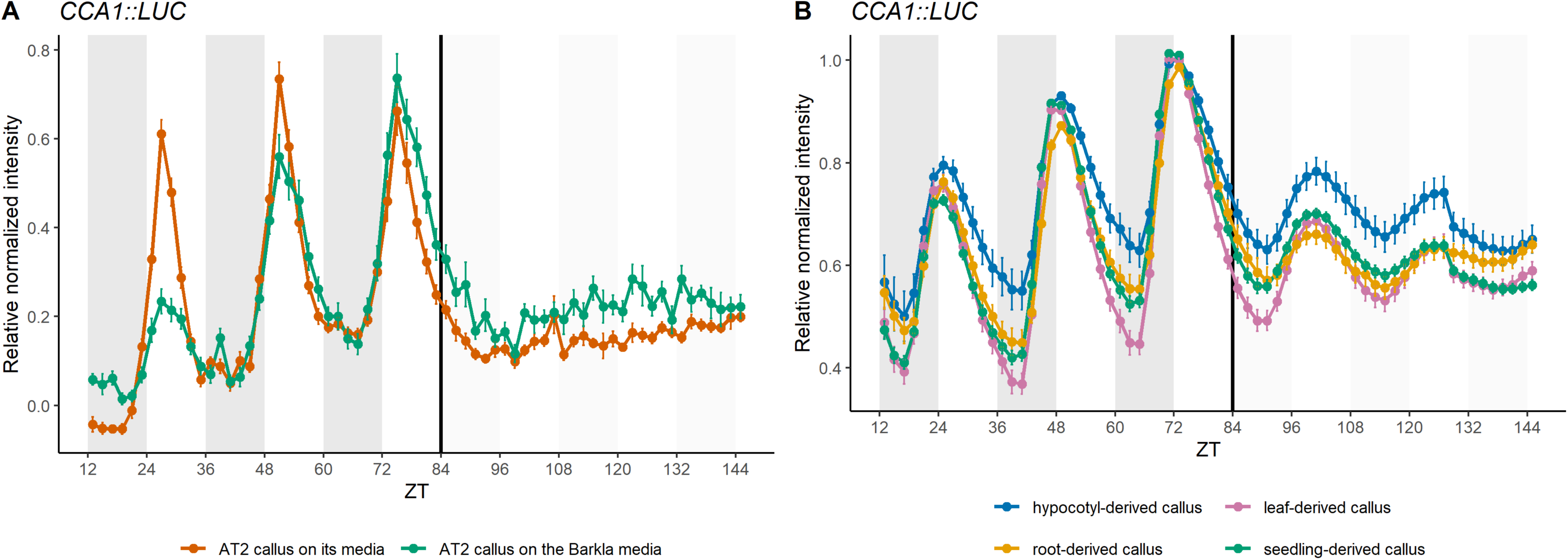
Expression of *CCA1::LUC* reporter is not affected by growth media or source tissue. (A) Bioluminescence assay of *CCA1::LUC* expression in the AT2 callus grown on the selective media (n=16) and the Barkla media (n = 16) under LD for 72 hours and subsequent LL for 60 hours. Data are mean ± SEM. (B) Bioluminescence assay of *CCA1::LUC* expression in calli derived from hypocotyls (n = 12), leaves (n = 12), roots (n = 12), and seedlings (n = 18) on the Barkla media under LD for 72 hours and subsequent LL for 60 hours. Data are mean ± SEM. n means the number of individual callus groups.

**Table 5.** Period lengths of the *CCA1::LUC* reporter are represented as mean ± SD for the AT2 callus grown on its media and the Barkla media under LD for 60 hours. n refers to a number of individual callus groups. Two-sample Wilcoxon rank-sum test was used to compare the means between own media and Barkla media; * = significant at P ≤ 0.05.

| Gene | AT2 media | Barkla media |
| --- | --- | --- |
| *CCA1::LUC* | 24.47 ± 0.43 (n=18) | 24.35 ± 2.74 (n=30) * |

The AT2 cell suspension was initiated from Col-0 leaf tissues while the callus we compared it to was generated from whole seedlings which were composed of various tissue types. We wanted to evaluate if the original tissue source affects the circadian oscillation in the callus since different organs exhibit a variation in robustness and precision of circadian oscillations (Takahashi et al. 2015). We produced calli from hypocotyl, leaves, and roots using the Barkla media and measured *CCA1::LUC* expression under LD and LL along with the callus derived from seedlings. The calli from different tissues showed similar *CCA1::LUC* waveforms and periods in LD (Figure 4B and Table 6). In the subsequent LL, callus cells derived from various tissues also maintained their rhythmic oscillations with no significant difference in periods between LD and LL (Figure 4B and Table 6). This is consistent with prior reports that all callus cells form a meristematic state that is similar to developmental lateral root formation (Sugimoto, Jiao, and Meyerowitz 2010; Fan et al. 2012; Ikeuchi, Sugimoto, and Iwase 2013; Atta et al. 2009). Therefore, these results, consistent with prior literature, suggest that the loss of rhythmic gene expression in constant light in the AT2 calli is not due to the explants used to generate the callus.

**Table 6.** Period lengths of *CCA1::LUC* in calli derived from different tissues on the Barkla media under LD and LL for 60 hours are represented as mean ± SD. n refers to a number of individual callus groups. One-way ANOVA test was performed to compare the means between tissues in LD. P-value = 0.132, indicating that there is no significant difference (P > 0.05). Paired-sample Wilcoxon test was used to compare the means between LD and LL; ns = not significant (P > 0.05) and * = significant at P ≤ 0.05.

| Tissues | LD | LL |
| --- | --- | --- |
| hypocotyls | 24.05 ± 0.21 (n=12) | 24.00 ± 0.74 ns |
| leaves | 24.15 ± 0.11 (n=12) | 24.27 ± 0.34 ns |
| roots | 24.11 ± 0.24 (n=9) | 23.25 ± 1.58 ns |
| seedlings | 24.20 ± 0.12 (n=18) | 24.41 ± 0.30 * |

### 3.4 Polymorphisms and altered expression of circadian clock genes are observed in AT2 cells

Another possible mechanism for the observed loss of rhythmicity in LL conditions is the presence of genomic mutations in the AT2 cells acquired during their maintenance in the pluripotent state. Such mutations could change gene expression at transcription or protein levels. As the time-course luciferase imaging could only determine the expression of a few circadian-associated genes in the AT2 cells, we analyzed RNA-Seq data previously performed on the AT2 cell suspension lines (PRJNA412215 and PRJNA412233) to evaluate the expression of 15 circadian-associated genes. The available RNA-Seq data was from AT2 cell suspensions grown in LL and DD. As a baseline, we compared expression levels of the circadian-associated genes to RNA-Seq data from Col-0 seedlings grown in LL at ZT0 and ZT12 (Grinevich et al. 2019) (Supplementary Figure 4A). Most of these 15 circadian-associated genes peak in expression either near ZT0 or ZT12. Since there was no record of the time of day when the AT2 cell suspension was harvested, it is not possible to evaluate their expression patterns from this data alone. However, overall, the total counts of most clock genes were substantially lower in both LL and DD in the AT2 cell suspension than at either ZT0 or ZT12 in Col-0. Only *PRR5*, *TOC1*, and *GI* had similar levels in the cell suspension and Col-0 (Supplementary Figure 4A). The expression of *CCA1* and *ELF4* in the AT2 callus were 19 times and 5 times lower than the lowest expression of those genes in seedlings at AM and PM, respectively (Supplementary Figure 4A). In addition to the difference in phases we would expect with these circadian-associated genes, they are often highly expressed in RNA-Seq datasets. We calculated the percent rank of gene expression to determine the ranks of clock genes among expressed genes in Col-0 seedlings and AT2 cell suspension. In Col-0 seedlings, the core clock genes such as *CCA1*, *LHY*, *PRR7*, *PRR9*, *TOC1*, *LUX*, *ELF3*, and *ELF4* were ranked in the top 20% at their highest (Supplementary Figure 4B). Even at the time when they are expected to be at their trough of expression, many circadian-associated genes remain in the top 50% of gene expression by counts (Supplementary Figure 4B). However, in the AT2 cell suspension, those genes were ranked as very lowly expressed with many in the lowest 30% of expressed genes in both data sets (Supplementary Figure 4B). This suggests that the expression of clock genes was not robust in the AT2 cell suspension.

We used the RNA-Seq data from the AT2 cell suspension to identify SNPs in the clock genes listed in (Nakamichi 2011) (Figure 5A and Supplementary Table 3). We found the mutations identified in *CASEIN KINASE II BETA CHAIN 3 (CKB3)*, *LIGHT-REGULATED WD (LWD1), LOV KELCH PROTEIN 2 (LKP2)*, *GI*, and *ELF3*. SNPs in *CKB3* and *LWD1* occurred in the 3’ UTR while the SNP in *LKP2* resulted in no amino acid changes (Supplementary Table 3). There were two SNPs in GI, one causing no changes in amino acid and another leading to premature stop. The premature stop was at the amino acid position 1156 in the last exon which was close to the true stop codon (position 1174). In the literature, the *gi-5* mutant (Fowler et al. 1999) has a base deletion in the last exon, which alters the last eight amino acids and adds 27 more amino acids at the C-terminus. The *gi5* mutant has similar gene expression as wildtype plants and shows a mild alteration in phenotype (late flowering only in the long-day condition) (Fowler et al. 1999; Mishra and Panigrahi 2015).

**Figure 5.**
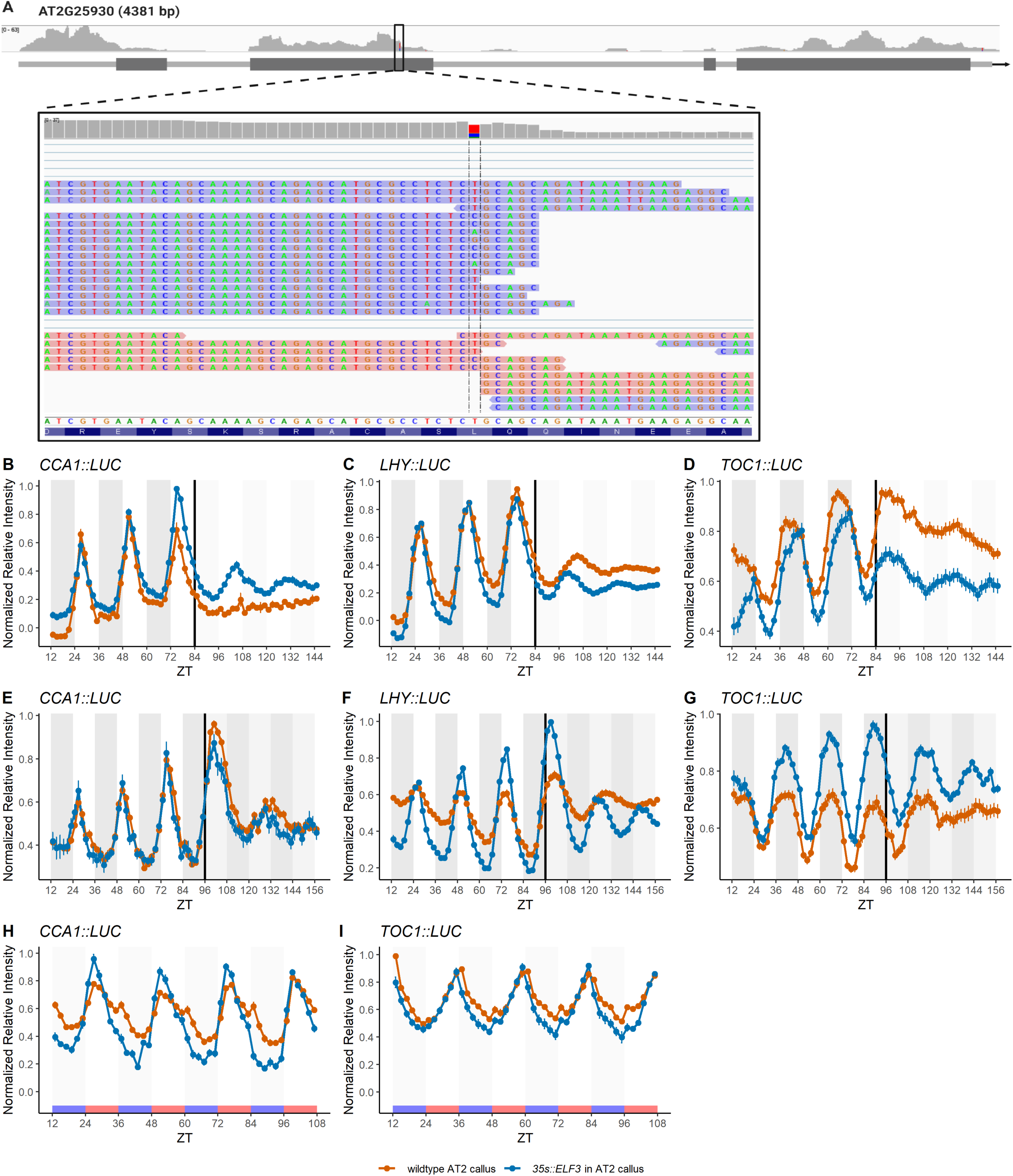
The overexpression of *ELF3* partially restored the circadian oscillations in the AT2 callus. (A) Location of the SNP on the second exon of the *ELF3* gene in the AT2 cell suspension. The sequences were visualized on IGV version 2.8.13. (B-G) Bioluminescence assay of *CCA1::LUC* (B and E), *LHY::LUC* (C and F), and *TOC1::LUC* (D and G) in the wildtype and *ELF3*- overexpressing AT2 calli under LD for 84 hours and subsequent LL for 96 hours (B-D) and LD for 96 hours and subsequent DD for 96 hours (E-G). Data are mean ± SEM. n = 18-35 individual wildtype callus groups and n = 10-30 individual *ELF3*-overexpressing callus groups. (H-I) Bioluminescence assay of *CCA1::LUC* (H) and *TOC1::LUC* (I) in the wildtype and *ELF3*- overexpressing AT2 calli under LLHC for 96 hours. Data are mean ± SEM. n = 18-36 individual wildtype callus groups and n = 12 individual *ELF3*-overexpressing callus groups.

An SNP (T to C) was also observed in the second exon of the *ELF3* gene resulting in a change in the amino acid position 291 (Figure 5A and Supplementary Table 3). CUG encoding leucine was changed to CCG encoding proline in the AT2 cell suspension (Figure 5A). The ELF3 protein is rich in serine, proline, and glutamine, and it does not share any domains that could be found in other protein families (Hicks, Albertson, and Wagner 2001). The ELF3 protein is divided into four regions based on amino acid richness: an acidic region (206-320), a proline-rich region (440-540), a threonine-rich region near the C terminus (636-652), and glutamine repeats or polyQ (544-585) in the region that has been recently predicted as a prion-like domain (430–609) (Hicks, Albertson, and Wagner 2001; Jung et al. 2020). The tertiary structure of ELF3 protein has not been constructed, so we cannot predict how this amino acid change would impact protein folding. We did sequence alignment to determine how conserved the leucine 291 was among plant species (Supplementary Figure 5, Supplementary Table 4, and Supplementary File 1). Leucine 291 was found mostly in dicot species (BrELF3a, VvELF3a, VvELF3, PtELF3a, and PtELF3b). GmELF3 had isoleucine and CpELF3 had phenylalanine instead of leucine. One rice ELF3 homolog had leucine like AtELF3, but other homologs in monocot species and lower plants (moss and spikemoss) had no conserved amino acids at this position and, in fact, many had a deletion of about 15 amino acids in the region surrounding Leucine 291. The *ELF3* gene from *Brachypodium distachyon* and *Setaria viridis*, is able to rescue defects in Arabidopsis *elf3* mutant even though there is sequence conservation at leucine 291 and nearby amino acids (Huang et al. 2017). This suggests that changes in protein sequences might not strongly affect the ELF3 expression or function.

With the RNA-Seq data, we are unable to detect mutations in the non-coding regulatory regions. However, the RNA-Seq indicated that the expression of circadian clock genes in the AT2 cells might be unusual compared to intact plants. We performed qRT-PCR to examine the expression of clock-associated genes at ZT0 and ZT12, dawn and dusk respectively, since the time of day the published RNA-Seq was performed is not known. We expect to see significant differences in most of the tested clock-associated genes at these two time points in both LD and LL. Indeed, in entrained LD conditions, we observed significant differences in expression between these two time points for *CCA1*, *LHY*, *TOC1*, *ELF4*, and *GI* (Supplementary Figure 3A, 3B, 3C, 3E, and 3G). The expression of *GI* in AT2 callus cells was significantly different between ZT0 and ZT12 in LD, but this difference diminished under LL similar to other clock genes with no SNPs (*CCA1*, *LHY*, *TOC1*, and *ELF4*) (Supplementary Figure 3A, 3B, 3C, 3E, and 3G). This suggests that the observed SNPs in *GI* might not affect *GI’s* expression. Only *ELF3* in the AT2 callus was not significantly cycling in both LD and LL.

For comparison, intact seedlings show a significant difference in expression of *ELF3* and all other tested circadian-associated genes even at two closer time points, ZT8 and ZT12 (Supplementary Figure 6). *ELF3*, as a member of the evening complex, increases in expression between ZT8 and ZT12 in seedlings, consistent with its peak expression at this time (Supplementary Figure 6D). However, in the AT2 cells, the induction *ELF3* is not observed between these two time points (Supplementary Figure 6D). In *elf3* knockout plants, circadian rhythms were abolished in the free-running conditions but maintained rhythmic in light/dark cycles (Hicks et al. 1996a; Reed et al. 2000a; Thines and Harmon 2010), similar to the pattern we observed in the AT2 callus cells. Therefore, based on the SNPs in the ELF3 coding sequence and the altered expression patterns, we hypothesized that either misexpression of *ELF3* or the mutation in the coding sequence could be the reason behind the altered circadian rhythm in the AT2 callus.

### 3.5 ELF3 overexpression recovered the circadian oscillation in the AT2 callus cells

To test the role of ELF3 in the disrupted circadian expression in the AT2 callus, we overexpressed *AtELF3* in the AT2 callus and measured *CCA1::LUC*, *LHY::LUC*, and *TOC1::LUC* under LD and free-running conditions (LL and DD) (Figure 5B-5G). The expression under LD in wildtype and *ELF3*-overexpressing calli were similar. The periods of *CCA1::LUC*, *LHY::LUC*, and *TOC1::LUC* in the *ELF3*-overexpressed callus were in the 24-hour range although the period length was significantly longer or shorter than the wildtype callus, suggesting that AT2 callus maintained the 24-hour LD rhythms with the *ELF3* overexpression (Figure 5B to 5G and Table 7 and 8). In constant light, the *ELF3*-overexpressing callus recovered the oscillations of *CCA1::LUC*, *LHY::LUC*, and *TOC1::LUC* (Figure 5B-5D). The period of *LHY::LUC* in the *ELF3*-overexpression callus (24.10 ± 1.04 h) was close to 24 h, but the periods of *CCA1::LUC* and *TOC1::LUC* were 27.05 h ± 2.25 and 28.55 h ± 1.62, respectively (Table 7). The reporter genes were rhythmically expressed in the AT2 callus under DD, and the overexpression of *ELF3* improved the oscillations of *CCA1::LUC, LHY:: LUC,* and *TOC1::LUC* (Figure 5E-5G). The period length of all three reporters shortened in the *ELF3*-overexpressing callus under DD (Table 8). We also tested whether the *ELF3*-overexpressing callus displayed the temperature-responsive peak in temperature cycles (Figure 5H and 5I). The temperature-responsive peak was reduced in the *ELF3*-overexpressing callus compared to wildtype (Figure 5H). However, *TOC1::LUC* exhibited a temperature-responsive peak in response to the shift from cool to warm temperatures (12°C to 23°C) in both wildtype and *ELF3* overexpression calli (Figure 5I). The results in the free-running conditions and temperature cycles indicate that proper expression of a wildtype ELF3 was able to partially recover normal oscillations in the AT2 callus.

**Table 7.** Period lengths of circadian associated reporters in wildtype and *ELF3* overexpression AT2 calli under LD and LL for 72 hours. Data are represented as mean ± SD. n refers to a number of individual callus groups or seedlings. Two-sample Wilcoxon rank-sum test was used to compare the means between wildtype and *ELF3* overexpression calli in LD. The significance levels are shown in the LD column; ns = not significant (P > 0.05), † = significant at P ≤ 0.05, and ‡ = significant at P ≤ 0.0001. Paired-sample Wilcoxon test was used to compare the means between LD and LL. The significance levels are shown in the LL column; ns = not significant (P > 0.05), * = significant at P ≤ 0.05, ** = significant at P ≤ 0.01, *** = significant at P ≤ 0.001, and **** significant at P ≤ 0.0001.

| Gene | Genotype | LD | LL |
| --- | --- | --- | --- |
| *CCA1::LUC* | wildtype | 23.80 ± 0.41 (n=18) ‡ | NA |
|  | *ELF3*-oex | 24.35 ± 0.29 (n=29) | 27.05 ± 2.25 **** |
| *LHY::LUC* | wildtype | 23.69 ± 0.22 (n=17) ‡ | NA |
|  | *ELF3*-oex | 24.07 ± 0.22 (n=16) | 24.10 ± 1.04 ns |
| *TOC1::LUC* | wildtype | 25.18 ± 0.62 (n=18) † | NA |
|  | *ELF3*-oex | 24.36 ± 0.87 (n=9) | 28.55 ± 1.62 ** |

**Table 8.** Period lengths of circadian-associated reporters in the AT2 wildtype and *ELF3* overexpressing calli under LD and DD for 72 hours. Data are represented as mean ± SD. n refers to a number of individual callus groups or seedlings. Two-sample Wilcoxon rank-sum test was used to compare the means between wildtype and *ELF3* overexpression calli in LD. The significance levels are shown in the LD column; ns = not significant (P > 0.05) and ‡ = significant at P ≤ 0.0001. Paired-sample Wilcoxon test was used to compare the means between LD and DD. The significance levels are shown in the DD column; ns = not significant (P > 0.05), * = significant at P ≤ 0.05, ** = significant at P ≤ 0.01, *** = significant at P ≤ 0.001, and **** significant at P ≤ 0.0001.

| Gene | Genotype | LD | DD |
| --- | --- | --- | --- |
| CCA1::LUC | wildtype | 24.12 ± 0.38 (n=24) ns | 30.52 ± 1.37 **** |
|  | ELF3-oex | 24.09 ± 0.72 (n=14) | 28.66 ± 2.90 ** |
| LHY::LUC | wildtype | 23.78 ± 0.25 (n=32) ‡ | 28.87 ± 1.30 **** |
|  | ELF3-oex | 24.01 ± 0.17 (n=31) | 25.01 ± 2.02 * |
| TOC1::LUC | wildtype | 24.39 ± 0.29 (n=13) ns | 29.27 ± 2.36 *** |
|  | ELF3-oex | 24.41 ± 0.25 (n=32) | 28.98 ± 1.04 **** |

## 4. Discussion

### 4.1 The AT2 callus system had altered circadian oscillations

Callus and cell suspension culture are valuable tools in plant basic research and industrial application (Efferth 2019). However, reports vary on if callus have a functional circadian clock ((Nakamichi et al. 2003, 2004; K. Lee, Park, and Seo 2016). Therefore, we investigated the oscillation of four circadian-associated genes *CCA1*, *LHY*, *TOC1*, and *FKF* in the AT2 callus under diel cycles and free-running conditions. We found that the expression of those genes was normal under LD. However, the rhythmic expression was lost in LL but maintained in DD albeit with lengthening periods. Callus needs sucrose as an external carbon source, and this could contribute to the maintenance of rhythmic expression under DD (Thorpe and Meier 1973). In Arabidopsis seedlings, the addition of sucrose to growth media results in a shortening period in constant light but helps maintain the robust rhythms in the constant dark condition (Knight, Thomson, and McWatters 2008; Dalchau et al. 2011; Haydon et al. 2013). In the AT2 callus, we observe a complete loss of period in constant light, suggesting that the sucrose in the media is not the only factor in the disrupted rhythmicity of the AT2 callus. Loss of rhythms in LL indicates that the AT2 callus has a disrupted circadian clock.

To evaluate if the loss of rhythms in AT2 callus cells was unique to this line or a result of the growth conditions, including media composition or callus source tissue, we generated fresh callus from seedlings. Unlike the AT2 callus, the seedling-generated callus from all tissue sources tested maintained oscillations in both LL and DD (Figure 4B, Table 4). These oscillations persisted on different media suggesting that the loss of rhythmicity in LL was unique to the AT2 callus and not the media composition. The observed rhythmicity in callus for these reporters is consistent with prior publications (W.-Y. Kim, Geng, and Somers 2003; Nakamichi et al. 2003, 2004; Xu, Xie, and McClung 2010; J. Sai and Johnson 1999). However, we observed shorter period lengths of *CCA1::Luc* in constant conditions in callus derived from all tissues than in intact seedlings. If this effect persists for other circadian reporters, this could be relevant for understanding how the cellular organization or identity contributes to circadian rhythms.

The loss of rhythm we observe in the AT2 callus has been previously observed in Arabidopsis cell lines. Nakamichi et al. 2003 observed rhythmic expression of the *PRRs* genes (*PRR1*/*TOC1*, *PRR5*, *PRR7*, and *PRR9*) in the T87 Arabidopsis cell line under LD and DD but not in LL (Nakamichi et al. 2003), a phenotype similar to what we observed with the AT2 callus. However, a year later, the same group reported that the T87 cell line had a functional clock because the expression of *CCA1::LUC* and *TOC1::LUC* was rhythmic under both LL and DD (Nakamichi et al. 2004). They suggested that the growth phase could affect the observation of circadian rhythms in the cell suspension. They used the callus grown on agar plates for over two weeks in the first publication but used fresher callus (three days on agar plates) in the experiment where the cells showed rhythmic expression. To ensure that the growth phase was not an issue with the rhythmicity, we grew the AT2 callus on the agar plates for 7-12 days before imaging to ensure that the AT2 callus and cell suspension were still in the exponential phase during this period (Supplementary Figure 7).

We evaluated if the response of the AT2 callus to temperature changes, another circadian input, was altered compared to intact seedlings. In the AT2 callus and callus derived from seedlings, the morning gene *CCA1* showed a small peak when the temperature decreased at dusk (22°C to 12°C), but this temperature-response peak did not occur in intact seedlings (Figure 3A). However, the evening gene *FKF* showed a similar temperature response when the temperature rose in the morning (12°C to 22°C) in both callus and seedlings (Figure 3D). To our knowledge, these temperature-responsive peaks to the daily temperature cycles have not been reported in intact plants so far (McWatters et al. 2000b; Somers et al. 1998; Salomé and McClung 2005). However, (Kusakina, Gould, and Hall 2014) showed that there was a second peak of *TOC1*::*LUC* occurring at the dark-to-light transition in seedlings grown in light/dark cycles at constant temperatures (12, 17, and 27°C), and the peak was enhanced as temperature increased (Kusakina, Gould, and Hall 2014). They observed the expression of *CCA1*::*LUC*, *LHY*::*LUC*, and *CAB2*::*LUC*, and found a small peak showing up at the transition from light to dark under low temperatures (17 and 12°C) (Kusakina, Gould, and Hall 2014). In our results, we found the second peak at the transition of temperature in both LD and LL, indicating that these peaks are specifically temperature-responsive and persist in constant light conditions despite the lack of rhythmicity in constant light. This induction of the circadian clock genes by temperature responses can be studied in callus to improve our understanding of temperature signaling in plants.

We examined the ability of the AT2 callus to resume the rhythmic gene expression when returned to LD to evaluate if the circadian clock persists and the rhythm is masked under LL. We saw that when the callus was returned to the dark, *CCA1* expression in the AT2 callus and seedlings decreased slightly and then began rising in the middle of the night to peak, as expected for an intact clock, at the dawn of the next light period (Figure 1C). The expression of *TOC1* in the AT2 callus increased right after the dark period. In seedlings, *TOC1* started increasing at 4 hours before the dark period, and darkness immediately shut down the *TOC1* expression (Figure 1F). This completely opposite response of *TOC1* expression to the dark after growth in LL between seedlings and the AT2 callus suggests that the AT2 callus could provide a novel means to examine circadian connections in the AT2 callus. The responsiveness of *CCA1* and *TOC1* expression to the dark after growth in LL suggests that either LL is masking an underlying circadian rhythm or that transition from light to dark serves as an input to the circadian oscillator in the AT2 callus. The experiments performed here cannot distinguish between these two possibilities.

### 4.2 Callus induction media and types of explant did not alter circadian oscillation

In our attempt to uncover the cause of the loss of rhythmicity in the AT2 callus, we discovered that the media composition and the tissue source of the callus did not significantly affect the rhythmic expression of the core circadian genes in freshly-induced callus. The disrupted rhythms were unique to the AT2 callus. Callus induced from seedlings had circadian expression similar to seedlings. The callus induction media contains auxin and cytokinin, which are known to affect circadian rhythms (Michael F. Covington and Harmer 2007; Zheng et al. 2006; Hanano et al. 2006). Auxin decreases the amplitude of the circadian rhythm while cytokinin delays the phase in Arabidopsis seedlings (Michael F. Covington and Harmer 2007; Zheng et al. 2006; Hanano et al. 2006). We found that the callus induced by two different concentrations of 2,4-D and BA exhibited a similar rhythmic expression pattern, indicating that this difference in hormones did not contribute to the loss of circadian oscillation in the AT2 callus under LL (Figure 4).

We also found that calli induced from hypocotyls, leaves, and roots showed a similar expression pattern compared to callus from whole seedlings, suggesting that initial tissues did not affect the circadian rhythms in the callus (Figure 4). Several studies indicated that the circadian clock function is tissue- and cell-specific (Thain et al. 2002; Takahashi et al. 2015; Endo et al. 2014; Román et al. 2020; Gould et al. 2018; James et al. 2008; Greenwood et al. 2019). At the tissue level, the root clock is less robust and precise than the shoot clock (Thain et al. 2002; Takahashi et al. 2015; Chen et al. 2020). Aerial and belowground tissues are composed of different cell types and experience different environments which result in altered oscillations (Sorkin and Nusinow 2021). Signaling molecules such as sucrose and ELF4 transmit circadian information to synchronize the clocks across tissues (Chen et al. 2020; James et al. 2008). It is possible that explants are transformed to callus via the same molecular mechanism regardless of types and concentrations of auxin and cytokinin or tissue source. As callus is considered a unique type of plant tissue based on gene expression profiles (Tanurdzic et al. 2008; He et al. 2012; Du et al. 2019; Shim et al. 2020; K. Lee, Park, and Seo 2018), cellular reprogramming during callus formation could resynchronize the clocks between cells, making callus cells exhibit similar circadian rhythms regardless of the origin of explants. Importantly, the seedling-like rhythms obtained in callus, independent of callus source or media, indicate that freshly-derived callus can be a useful tool for studying circadian regulation.

### 4.3 ELF3 recovered the circadian defect in the AT2 callus

We hypothesized that the difference in rhythmic gene expression between the freshly-derived callus and the AT2 callus could be due to one or more mutations accumulated in the AT2 cell suspension. We examined the RNA-Seq data and found that the expressions of clock genes were relatively low compared to intact plants in LD and free-running conditions. We found that, unlike in seedlings, the expression of *ELF3* was not significantly different between dawn and dusk in both LD and LL, and there was one SNP in the second exon of *ELF3* which led to leucine-to-proline mutation at position 291. *ELF3* is a key circadian component involved in light and temperature signalings, regulation of flowering, and thermomorphogenesis (Box et al. 2015; McWatters et al. 2000b; Nusinow et al. 2011b; Zagotta et al. 1996; Jung et al. 2020).

The protein sequence alignment indicates that leucine 291 is not conserved among species (Supplementary figure 5). Monocot ELF3 proteins lack leucine 291 and nearby amino acids but they can complement the functions of ELF3 in Arabidopsis *elf3* knockout (Huang et al. 2017). This suggested that changes in leucine 291 may not strongly impact the functions of ELF3. Although we did not assess whether the SNP affects protein function as we do not know the folding of ELF3 protein. Previous ELF3 studies showed that *elf3* knockout exhibited arrhythmic *CAB2* and *CCR2* in LL but not in DD (Hicks et al. 1996b; M. F. Covington et al. 2001b; Reed et al. 2000b). But Thines and Harmon found that the *elf3* knockout had no rhythmic *TOC1*, *CCR2*, and *FKF* expression in both LL and DD (Thines and Harmon 2010). The authors commented that the rhythmic expression in the previous publication could be driven by the last light cycles prior to free-running conditions (Thines and Harmon 2010).

Overexpressing ELF3 in the AT2 callus restored circadian expression in the free-running conditions and temperature cycles resulting in rhythms similar to that in seedlings (Figure 5). Overexpression of ELF3 in seedlings resulted in robust circadian rhythms DD but slightly lengthening periods in LL (M. F. Covington et al. 2001b). Despite a reduced minor peak of *CCA1*::*LUC* in the *ELF3*-overexpressed calli under temperature cycles, we found that the minor peak of *TOC1*::*LUC* still persisted suggesting that some subtle differences remain between the *ELF3-*overexpressing AT2 cells and seedling circadian gene express.

Overall, this study shows that the expression of circadian clock genes in the AT2 callus is driven by light and temperature signals but AT2 callus exhibits no circadian oscillations in constant light conditions. The overexpression of ELF3 in the AT2 callus restores rhythmic expression in LL. Rhythmic expression of clock genes was robust in freshly made calli in both LD and LL despite induction media composition and explant source tissue. This suggests that callus could be useful in clock studies, but caution should be employed to ensure that the callus or cell suspension cultures do not lose their rhythmic expression.

## Acknowledgements

We would like to thank Linda Hanley-Bowdoin and Wei Shen, North Carolina State University, for providing the AT2 cell suspension and cell maintenance and transformation protocols. We appreciate comments and suggestions from Jose Pruneda-Paz, UC San Diego, on this manuscript. KL was supported by the Development and Promotion of Science and Technology Talents Project (DPST), Thailand. Additional support was provided by USDA National Institute of Food and Agriculture Project 1002035.

## Authors’ Contributions

C.J.D. conceptualized the experiments. K.L. grew callus and plants for luciferase imaging, analyzed bioluminescence data and RNA-Seq data, and performed qRT-PCR. J.S.D. identified SNPs from the RNA-Seq data. All authors participated in writing the manuscript.

---

**Supplementary Figure 1 Gentamicin did not interfere with the clock gene expression in the AT2 callus under LD**

Bioluminescence assay of *CCA1::LUC* (A), *LHY::LUC* (B), *TOC1::LUC* (C), and *FKF1::LUC* (D) in the AT2 callus grown on gentamicin-containing media and regular media under LD for 60 hours. Data are mean ± SEM. n = 17-18 individual callus groups on gentamicin-containing media and n = 13-36 individual groups on regular media.

**Supplementary Figure 2 The AT2 callus had an altered circadian oscillation**

Bioluminescence assay of *LHY::LUC* (A-C) and *FKF1::LUC* (D-F) in the AT2 callus and seedlings under LD for 72 hours and subsequent LL for 60 hours (A and D), LD for 84 hours and subsequent DD for 60 hours (B and E), and LL for 48 hours and subsequent LD for 58 hours (C and F). Data are mean ± SEM. n = 14-36 individual callus groups and n = 10-41 seedlings.

**Supplementary Figure 3 Most clock-associated reporters lost their robust daily variation in expression in the AT2 callus under LL**

Relative normalized expression (ΔΔCq) of *CCA1* (A), *LHY* (B), *TOC1* (C), *ELF3* (D), *ELF4* (E), *LUX* (F), and *GI* (G) in the AT2 callus at ZT0 and ZT12 under LD (blue) and LL (green). Data are an average of ΔΔCq relative to zero ± SD and n = 3-4. The expression was normalized by *IPP2* and external RNA spikes. Two-sample t-test was used to compare the means between ZT0 and ZT12: ns = not significant (P> 0.05), * = significant at P ≤ 0.05, ** = significant at P ≤ 0.01, *** = significant at P ≤ 0.001, and **** = significant at P ≤ 0.0001.

**Supplementary Figure 4 Circadian-associated genes showed relatively low expression in the AT2 cell suspension.**

Heat maps of the log_10_-transformed normalized read counts (A) and the percent rank (B) of gene expression in the AT2 cell suspension grown in LL and DD (‘AT2_LL’ and ‘AT2_DD’) compared to Col-0 seedlings at dawn and dusk in LL (‘seedlings_LL_AM’ and ‘seedlings_LL_PM columns’).

**Supplementary Figure 5 ELF3 protein sequence alignment indicates that Leucine 291 is conserved among dicot species but not in monocot groups.**

ELF3 protein sequences from different plant species were retrieved from Phytozome version 13 and aligned by Clustal Omega version 1.2.2 on Geneious Prime® 2021.2.2 with default options. Highlights are amino acids that agree with the consensus sequence. The full alignment is shown in Supplementary Information File 1. Transcript identifiers are shown in Supplementary Table 4.

**Supplementary Figure 6 Variation in expression of circadian-associated genes in the AT2 cell suspension was lower than in intact plants.**

Relative normalized expression (ΔΔCq) of *CCA1* (A), *LHY* (B), *TOC1* (C), *ELF3* (D), and *ELF4* (E) in Col-0 seedlings and the AT2 cell suspension harvested at ZT8 and ZT12 under LD. Data are an average of ΔΔCq relative to zero ± SD and n = 3-4. The expression was normalized by *ACT2*, *TUB2/3*, *IPP2* and external RNA spikes. Two-sample t-test was used to compare the means between ZT8 and ZT12; ns = not significant (P > 0.05), * = significant at P ≤ 0.05, ** = significant at P ≤ 0.01, *** = significant at P ≤ 0.001, and **** = significant at P ≤ 0.0001.

**Supplementary Figure 7. *Agrobacterium*-transformed AT2 cell suspension grew slower than the wildtype AT2 cells.**

Growth curves of (A) AT2 cells (n = 4) and (B) *Agrobacterium*-transformed AT2 cell suspension containing *CCA1::LUC*, *LHY::LUC*, *TOC1::LUC*, and *FKF1::LUC* (n = 2 of each construct). The red horizontal line indicated 50% of the total cell volume. Data are mean ± SEM.

**Supplementary Table 1. Primers used for qRT-PCR**. Primer name indicates the transcript targets and if the primer is forward (f) or reverse (r).

**Supplementary Table 2. Periods and amplitudes of circadian-associated reporters in seedlings under LD and subsequent LL for 60 hours.** Data are represented as mean ± SD. n refers to a number of individual seedlings. Paired-sample Wilcoxon test was used to compare the means between LD and LL; ns = not significant (P > 0.05), * = significant at P ≤ 0.05, and ** = significant at P ≤ 0.01

**Supplementary Table 3. SNPs identified in AT2 cells from RNA-Seq data.** The gene list is from Nakamichi, N. (2011). The RNA-Seq data was downloaded from NCBI SRA (PRJNA412215 and PRJNA412233).

**Supplementary Table 4. Gene Identifiers for ELF3 homologs from different plant species.**

Gene identifiers used for ELF3 homologs in Supplementary Figure 5 and Supplementary File 1.

**Supplementary File 1. ELF3 alignment across species.** Alignment of ELF3 proteins across several plant species. Arabidopsis ELF3 is shown in red. Leucine 291 is emphasized by increased font size and is underlined.

