## Supplementary figures and images for "Arabidopsis cell suspension culture that lacks circadian rhythms can be recovered by constitutive ELF3 expression"

### Supplementary Figure 1

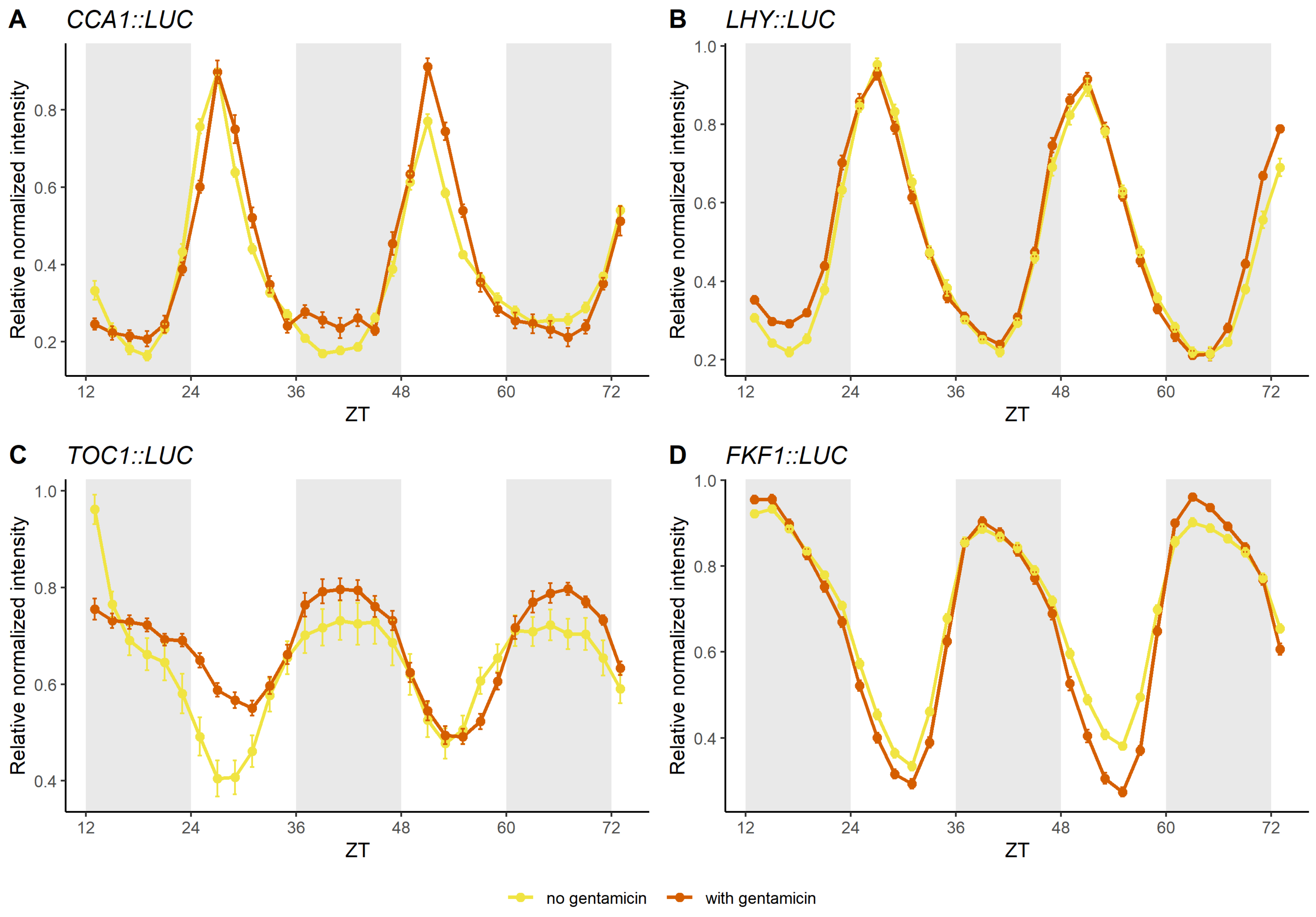

### Supplementary Figure 2

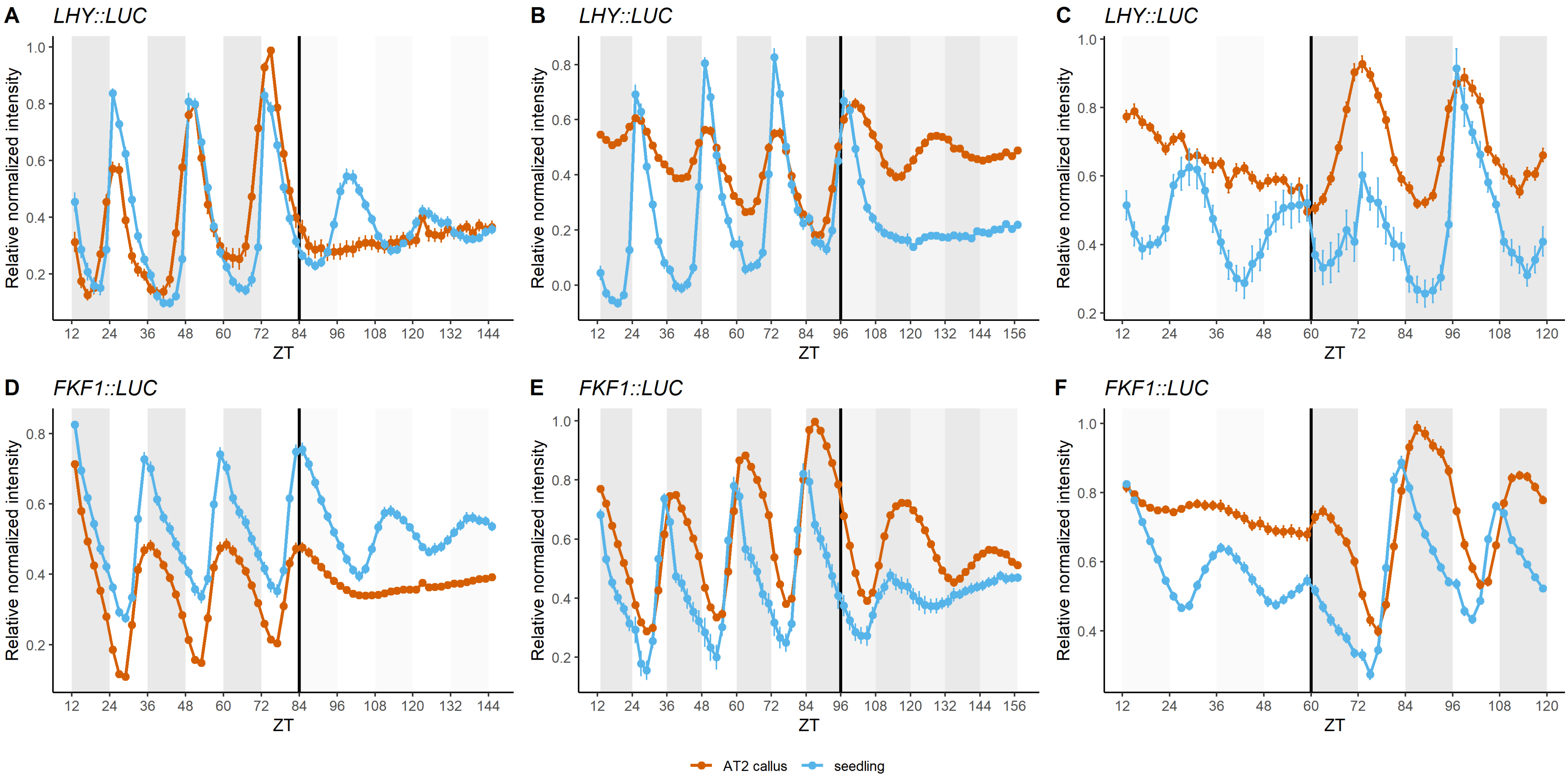

### Supplementary Figure 3

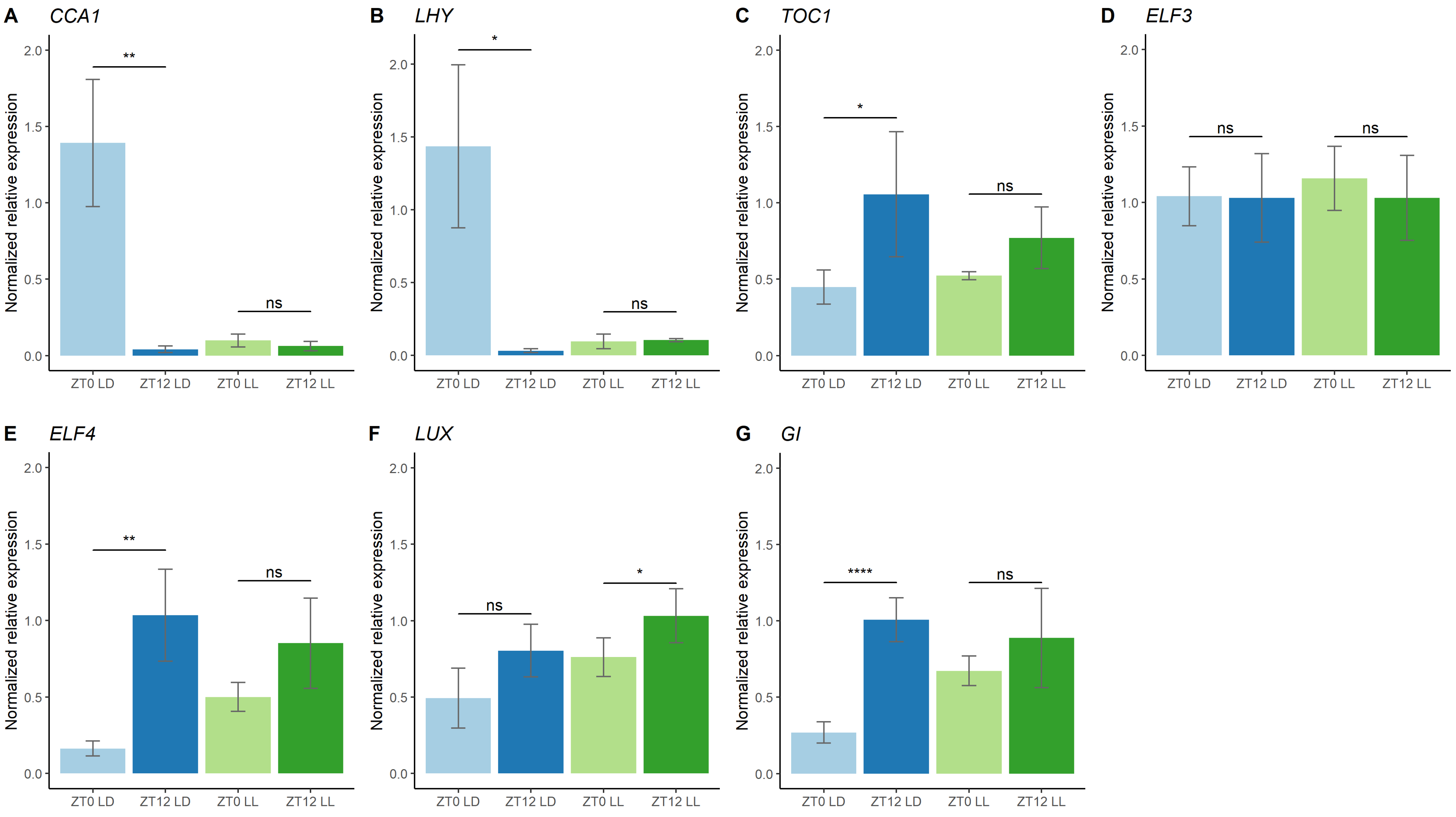

### Supplementary Figure 4

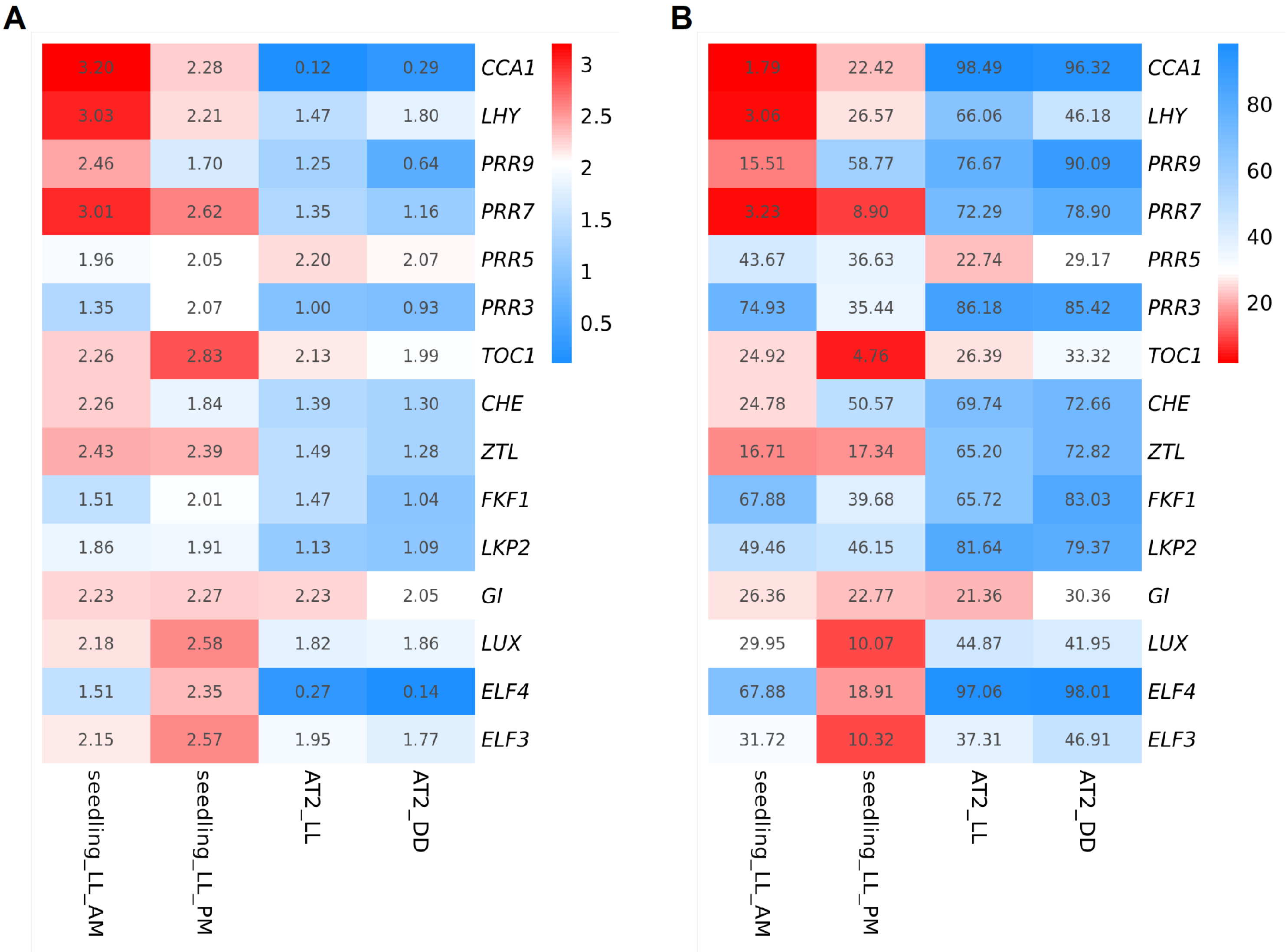

### Supplementary Figure 5

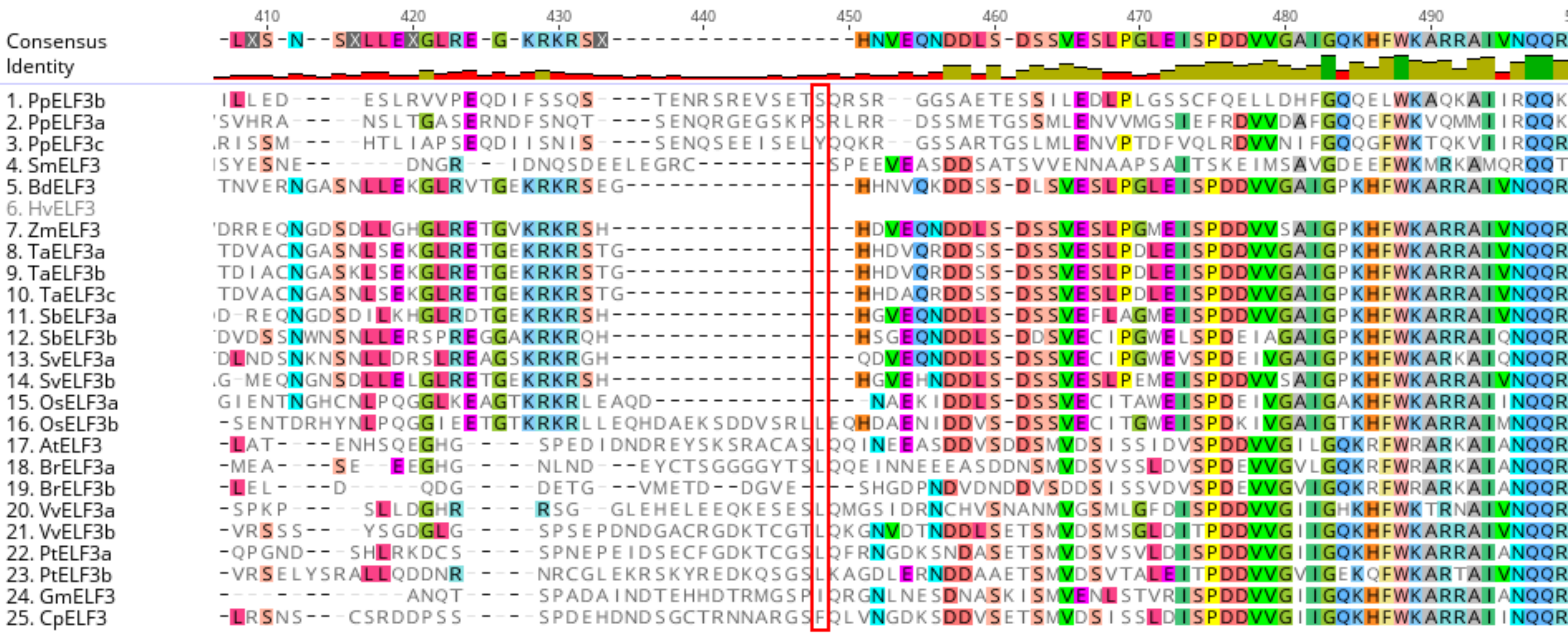

### Supplementary Figure 6

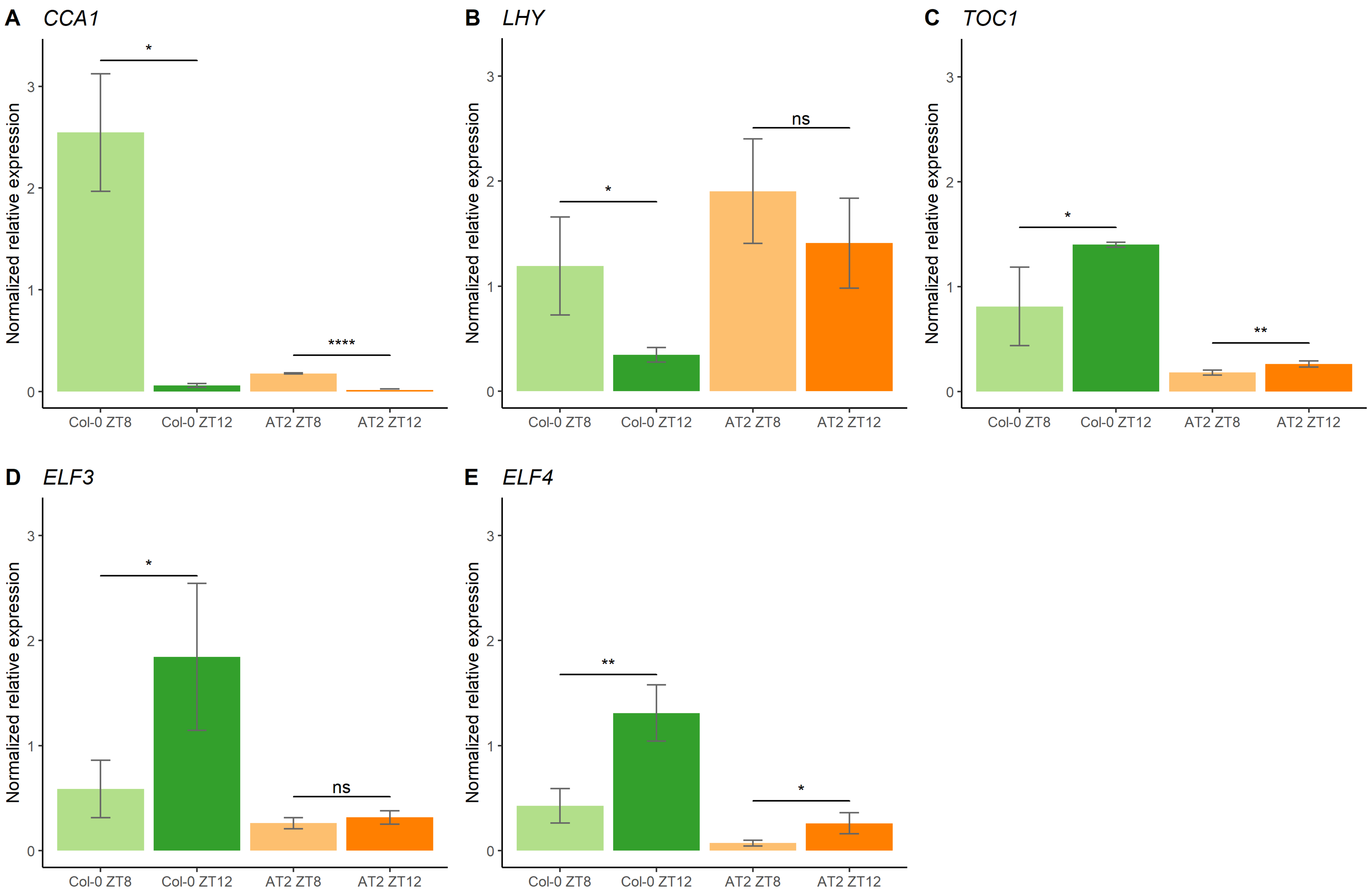
