## Supplementary Tables for "Arabidopsis cell suspension culture that lacks circadian rhythms can be recovered by constitutive ELF3 expression"

| **Primer name** | **Sequence in the 5’-to-3’ direction** |
| --- | --- |
| CCA1_f | GTCTGACGAGGGTCGAATTG |
| CCA1_r | GTTGTTGTTGTTCTTCCTCTCTG |
| LHY_f | ACCAACGAAACAGGTAAGTGGCG |
| LHY_r | GTGCACAGTAGTACCATCGTTACC |
| TOC1_f | AGAAATCCAGCGCAATTTTCTTCAG |
| TOC1_r | GACCGTTAGTTCTAAGGACAG |
| ELF3_f | GCACAGACTGATTAAGGTTCAAAAAC |
| ELF3_r | CTTCACTGGATAGCTTTTAGCAG |
| ELF4_f | TGTCGTTGACTTGTTGAATCAGTG |
| ELF4_r | CGATGTGGGAGAATCTTGAC |
| LUX_f | GGAAATGGTGCGTTTGAGTC |
| LUX_r | TTCTCATTTGCGCTTCCACC |
| GI_f | CTCAACTTGAGCTACTTGAAGCC |
| GI_r | GGTAGACGACACTTCAATAGATTG |
| PRR5_f | GAGAAAAAGGTTCGTTACGAGAG |
| PRR5_r | AGCTTGTGTGGATTGGACTTG |
| IPP2_f | GTATGAGTTGCTTCTCCAGCAAAG |
| IPP2_r | GAGGATGGCTGCAACAAGTGT |
| Artificial spike 2_f | CCAGAATGGCTTCCAGCTTTA |
| Artificial spike 2_r | GGAACAAGACCGCAAGGAATTA |
| ACT2_f | CGTACAACCGGTATTGTGCTGG |
| ACT2_r | AGCAAGGTCAAGACGGAGGATG |
| TUB2/3_f | CCAGCTTTGGTGATTTGAAC |
| TUB2/3_r | CAAGCTTTCGGAGGTCAGAG |

**Supplementary Table 1. Primers used for qRT-PCR.** Primer name indicates the transcript targets and if the primer is forward (f) or reverse (r).

| Gene | Period (hours) | | Amplitude | |
| --- | --- | --- | --- | --- |
|  | LD | LL | LD | LL |
| *CCA1::LUC* | 24.29 ± 0.38 (n=10) | 24.97 ± 0.77 * | 0.2500 ± 0.0327 (n=10) | 0.1270 ± 0.0298 ** |
| *LHY::LUC* | 24.01 ± 0.25 (n=7) | 25.02 ± 0.57 * | 0.3071 ± 0.0403 (n=7) | 0.2243 ± 0.0503 * |
| *TOC1::LUC* | 24.14 ± 0.26 (n=7) | 26.85 ± 1.18 * | 0.3000 ± 0.0224 (n=7) | 0.2014 ± 0.0313 * |
| *FKF1::LUC* | 24.26 ± 0.65 (n=7) | 27.06 ± 0.98 * | 0.1843 ± 0.0190 (n=7) | 0.1143 ± 0.0199 * |

**Supplementary Table 2. Periods and amplitudes of circadian-associated reporters in seedlings under LD and subsequent LL for 60 hours.** Data are represented as mean ± SD. n refers to a number of individual seedlings. Paired-sample Wilcoxon test was used to compare the means between LD and LL; ns = not significant (P > 0.05), * = significant at P ≤ 0.05, and ** = significant at P ≤ 0.01

| **Name** | **AGI** | **coordinate** | **Original base (%occurrence) -> SNP (% occurrence)** | **Location and outcome** | |
| --- | --- | --- | --- | --- | --- |
| CKB3 | AT3G60250 (gene on minus strand) | 3:22270370 | A (55%) -> G (25%)  A (55%) -> C (3%) | 3’ UTR | |
| LWD1 | AT1G12910 (gene on - strand) | 1:4395028 | A (66%) -> G (17%)  A (66%) -> C (10%) | 3’ UTR | |
| LKP2 | AT2G18915 (gene on - stand) | 2:8195698 | G (55%) -> A (27%)  G (55%) -> T (18%) | Second exon; (GGC) Glycine 310 -> (GGT) Glycine  Exon #9 CGG (Arg) -> AGG (Arg) | |
| GI | AT1G22770 | 1:8067391  1:8064843 | C (60%) -> T (40%)  C (72%) -> A (13%) | Last exon (#13): CAG (Gln) 1156 -> TAG (stop) | |
| ELF3 | AT2G25930 |  | T (59%) -> C (27%) | Second exon: (CTG) Leucine 291 -> (CCG) Proline | |
| Genes with no detected SNPs | | | | | |
| **Name** | **AGI** | **Name** | **AGI** | **Name** | **AGI** |
| CCA1 | AT2G46830 | PRR9 | AT2G46790 | PRMT5 | AT4G31120 |
| LHY | AT1G01060 | PRR7 | AT5G02810 | FKF1 | AT1G68050 |
| CKB4 | AT2G44680 | PRR5 | AT5G24470 | ZTL | AT5G57360 |
| CHE | AT5G08330 | PRR3 | AT5G60100 | JMJD5 | AT3G20810 |
| LUX | AT3G46640 | TOC1 | AT5G61380 | MYB3R2 | AT4G00540 |
| ELF4 | AT2G40080 | FIO1 | AT2G21070 | bHLH69 | AT4G30980 |
| LWD2 | AT3G26640 | TIC | AT3G22380 | TEJ | AT2G31870 |

**Supplementary Table 3 SNPs identified in AT2 cells from RNA-Seq data.** The gene list is from Nakamichi, N. (2011). RNA-Seq data is from

| Species | Gene name | Transcript identifier |
| --- | --- | --- |
| Moss (*Physcomitrium patens*) | PpELF3a | Pp3c1_12790V3.1 |
|  | PpELF3b | Pp3c11_14750V3.1 |
|  | PpELF3c | Pp3c7_10610V3.1 |
| Spikemoss (*Selaginella moellendorffii*) | SmELF3 | SmELF3415241 |
| Stiff brome (*Brachypodium distachyon*) | BdELF3 | Bradi2g14290.1 |
| Barley (*Hordeum vulgare*) | HvELF3 | HORVU1Hr1G094980.1 |
| Maize (*Zea mays*) | ZmELF3a | Zm00001d039156_T001 |
|  | ZmELF3b | Zm00001d044232_T001 |
| Bread wheat (*Triticum aestivum*) | TaELF3a | Traes_1BL_B95F8C666.1 |
|  | TaELF3b | Traes_1AL_52C5531A4.1 |
|  | TaELF3c | Traes_1DL_96D83DE2D.1 |
| Sorghum (*Sorghum bicolor*) | SbELF3a | Sobic.009G257300.2 |
|  | SbELF3b | Sobic.003G191700.1 |
| Green foxtail (*Setaria viridis*) | SvELF3a | Sevir.5G206400.1 |
|  | SvELF3b | Sevir.3G123200.1 |
| Rice (*Oryza sativa*) | OsELF3a | LOC_Os01g38530.1 |
|  | OsELF3b | LOC_Os06g05060.1 |
| Thale cress (Arabidopsis thaliana) | AtELF3 | AT2G25930.1 |
| Field mustard (Brassica rapa) | BrELF3a | Brara.I04408.1 |
|  | BrELF3b | Brara.D01576.1 |
| Wine grape (Vitis vinifera) | VvELF3a | VIT_209s0002g02680.2 |
|  | VvELF3b | VIT_204s0008g00660.1 |
| Black cottonwood (*Populus trichocarpa*) | PtELF3a | Potri.006G233800.1 |
|  | PtELF3b | Potri.003G045000.2 |
| Soybean (*Glycine max*) | GmELF3 | Glyma.04G050200.1 |
| Papaya (*Carica papaya*) | CpELF3 | evm.model.supercontig_78.11 |

**Supplementary Table 4 Gene Identifiers for ELF3 homologs from different plant species.** Gene identifiers used for ELF3 homologs in Supplementary Figure 6 and Supplementary File 1.

**Supplementary File 1** ELF3 protein sequence alignment using Clustal Omega version 1.2.2 on Geneious Prime® 2021.2.2 with default options. Dots indicate the position in which the amino acid agrees with the amino acid in the consensus sequence. Dashes indicate gaps.

[Sequences are in the docx file ‘ELF3_protein_alignment.docx’]
