## Supplementary File 1 for "Arabidopsis cell suspension culture that lacks circadian rhythms can be recovered by constitutive ELF3 expression"

Consensus MSFRNXMMKR-X----GKD-EDKVMGPLFPRLHVNDTX-KXGGPRAPPRNKMALYEQLSV	53

PpELF3b	-------..GAKEGKADGIKG--AGPA.......AE.K--KA...G............TI	49

PpELF3a	-------...EKGERSDGGKQQEPEIA.......AE.K--KA...G............T. 51

PpELF3c	.....IKT.GSMEGKADGSKQQE.GAA.......AE.K--KA...G........S...TI	58

SmELF3	----------------ME.KNAPPG.A.......KE.K--NA................TI	42

BdELF3	-------.R.GGGGPAPAGK.................L--K................F.. 51

HvELF3	------------------------------------------------------------

ZmELF3	-------.R.GAT--KDDAAP................L--K................F.. 49

TaELF3a	-------.R.AGG---KDGG.................TL.G................F.. 50

TaELF3b	-------.R.AGG---KDGG.................TL.G................F.. 50

TaELF3c	------------------------------------------------------------

SbELF3a	-------.R.GGA--KDDAAP................L--K................F.. 49

SbELF3b	------.TRGGGGGGD-------GK.........S.AG-.G.................T. 46

SvELF3a	------.TRGGAAGGG.REEQG............S.AG-.G................FT. 53

SvELF3b	-------.R-RGA--GKDEAP................V--K................F.. 48

OsELF3a	-------.RGGGGGGKEVEERG..............AA-.G................FT. 52

OsELF3b	-----MATRGGGGGGG..EAKG..............AA-.G................FT. 54

AtELF3	-------...------...-.E.ILE.M........AD-.-..................I	44

BrELF3a	-------...------..E-.E.RLE.M......K.AE-.-..................I	44

BrELF3b	-------...------..EE.E.KLE.M........AD-.G...............H.TI	46

VvELF3a	-------..K------...-.G...S..........AE-.-................... 44

VvELF3b	-------...------...-.E.I...M.........E-.-................... 44

PtELF3a	-------...------...-Y..I...M.........D-.-..................I	44

PtELF3b	------------------------------------------------------------

GmELF3	-------...------...-DE.....M.........E-.-................F.I	44

CpELF3	-------...------...-DE.ITE...........D-.-..................I	44

Consensus PSHRFSASXA-P------------------XXP--XSSLVPSAS---XSQVGGYDRPL-F	88

PpELF3b	....YKM.PL-.----------LPSSGRVMST.SMTTP.F-----NPQV.ASS..SR.PY	93

PpELF3a	...K.KP.PM-.----------LPPNGRTIQS.SMSTPIY-----LPQMP..S.ESMYSY	95

PpELF3c	...K.KP.PI-A----------LPSSGRNTKY.SMCTPTY-----LPQV.T.S..SMYS. 102

SmELF3	.....QQQQSDGAPPQ---------------------PF..QVCKFEPMISPALN.GY-Y	80

BdELF3	..Q..TPHR.S-------------------------..ALS...---PG.I..S....-. 82

HvELF3	------------------------------------------------------------

ZmELF3	....Y..AVP-------PAPSPAPPWGAQRPAS-----A...T.---A.....G...I-. 93

TaELF3a	..Q..A.NA.NTA-----------PAAAHRP.XXXSYAA.S...---AG.I..I....-. 95

TaELF3b	..Q..A.NA.NTA-----------PAAAHRPA--ASYAA.S...---AG.I..I....-. 93

TaELF3c	-------------------------------------------------RI..I....-. 10

SbELF3a	....Y..AA.A.PPPSAPAPSPAPPWRAQRPV.--------AT.---A.....S....-. 97

SbELF3b	..N...SPAPA.ARAA-------------GAKA----.....T.---AA..Y....T.-. 85

SvELF3a	..N...S.A.STRA----------------AGG----...T.T.---A...YS.....-. 89

SvELF3b	.......AA.A.A-----APAPAPPWHAHRPA.GAAT.A...T.---A..A..S....-. 99

OsELF3a	......GGGGALASAR-------------GSLAR----STSA..---Q...Y.C.M..-. 91

OsELF3b	......GGGGGGGVG--------------GSPAHST.AASQ.Q.---Q...Y.R.SS.-. 96

AtELF3	..Q..GDHGTMNS-----------------R-SNNT.T..HPGP---S..PC.VE.N.-S	82

BrELF3a	..Q..TS.DRVGG-----------------SL.RNT.P.L.PGPSSN-------------	74

BrELF3b	.....TDHHS--------------------SS.RHTNT.F.PPPPVPSN.PC.VE.N.-T	85

VvELF3a	.KVPSRSTVIL.------------------LL.NNSG......P---ST.GS.HE.NV-V	82

VvELF3b	..Q.CN-PGVM.------------------LKSNNA.N...P..---S..GS.HE.GV-. 81

PtELF3a	..Q..N-PDVL.------------------RNSSNT.D.A.TG.---S..GS.LE.HF-P	81

PtELF3b	------------------------------------------------------------

GmELF3	..Q..N-.GVL.------------------LN.NIS.NT..P..---S.LRTVPE.NC-V	81

CpELF3	..Q.Y.-PGVLA------------------LN.SNTN....PG.---ST.GH.IE.N.-. 81

Consensus PPFCVP-SN-EPVRS-SEKMNSNSSGRDNNST-----RXE-SGR---STQ-KSXD-XA-X	133

PpELF3b	M.MGYMG.TII.TMAFMPI..MGLY.VGAGADAGTGASAS-.SM--NASSS..TVVDGTT	150

PpELF3a	M..SYMKPAVV..MTCMPV...GQC.VEAK.S----------KI--T.SNSD.TVMDGAS	143

PpELF3c	V.MNYMN.GII.TMACVPM...GLYSV.SR.S----------LN--T-CSS..TVVDGSS	149

SmELF3	F.YYMVP.P-VAFQP-MNVVLGQH.QSVA.E-----------TEQRELQ.-----VPGDG	122

BdELF3	.S....-..-..A..-..HI.T..N...G.A.-----.V.-...--H...L..K.TY.-A	130

HvELF3	------------------------------------------------------------

ZmELF3	.L.R..-.T-.....-.DQT.A..N.QGA.G.-----IA.-...QRQ..HL..K.TN.-A	143

TaELF3a	.S....-..-....L-P.HIKT......GQA--------I-...--L...L..K.AY.-A	140

TaELF3b	.S....-..-....L-P.HIKT......GHA--------T-...--L..LL..K.AY.-A	138

TaELF3c	.S....-..-....L-P.HIKT......GHA--------T-...--L...L..K.AY.-A	55

SbELF3a	.S....-.T-.....-.DQ..A..N..AA.G.-----.A.-...--Q..HL..K.TN.-A	145

SbELF3b	Q..N..-..-..A..-...FKG.CINGQS...-----.R.-.L.--M.S.T..K.VC.-S	133

SvELF3a	Q..D..-..-G.AH.-...FKG..INGQS...-----.R.-...--MP..T.NN.VY.-S	137

SvELF3b	.S....-.T-.....-.DH..A..N..AG.A.-----.A.-...--L..HL..K.TN.-A	147

OsELF3a	E..N..-..-G.GQ.-V.......VN.QI.GS-----.KD-..M--L...P.GI.KYG-S	139

OsELF3b	Q..N..-..-R.GH.-T..I..DKINKKISGS-----.K.-L.M--L.S.T.GM.IY.-S	144

AtELF3	VQHLDS-.A-A-NQA-T..FV.QM.FME.VRS-----SA--QHDQR--------------	117

BrELF3a	-----Q-GT--------DNFVTQMPLME.VR.-----SSAQHDHQR--------------	101

BrELF3b	SQHLDS-.A--------SGHVTQM.SME.VT.-----LAHRR.DQR--------------	117

VvELF3a	T...YT-PT-S-AH.-D.NHHFH..SGV.M.F-----ELT-NLEQKSG.FYHIPNVTGQL	132

VvELF3b	F.HHIS-PS-T.THL-P..LHARH.DAVILN.-----PLA-QFEQR--------------	118

PtELF3a	Y.HY..-.P-T.TDV-A..YH.RQPDGR.LN.-----PVA-PLMQR--------------	118

PtELF3b	------------------------------------------------------------

GmELF3	Y.VHL.-PQ-R.IHR-A..C..RQ.EGT.L.A-----SL----EQR--------------	115

CpELF3	I.AHA.-PA-T.T.L-ADAF.GDQHNGA.V.G-----PTG-QQEQR--------------	118

Consensus --GS----AEC-S--R--------X--XXSSGKKLXNDDDFTVPSVFSSGXPPHSTQEXX	174

PpELF3b	ESQRVPWGG.AP.RAN----KSSDHAKHLAPS..GKE....A..TYS.ATQGSSQ.RPSI	206

PpELF3a	ESP.---LGRAVGRKS----ESLGHC---APTN.ESLG...A..TYS.ASQSSSQ.VLTT	193

PpELF3c	-EP.---VGGAVTQPP----KSFNHSKQH.PS..GSR..E.A..AHS.GTPSSLLQTQQD	201

SmELF3	--..NRQL.RIP.LHSR-------AGSGKTKTRAPRV..VAA..TYP.------------	161

BdELF3	--..T---...S.QR.E-------NSVKN......T...........C..V.......VV	178

HvELF3	------------------------------------------------------------

ZmELF3	--.PP---..G----------------NN.V....A..........LY..M....S..--	180

TaELF3a	--..T---...S.SQ.RDNNN---NSMKN......T...........C..AR.R.NH.EV	192

TaELF3b	--..T---...S.SQ.RDNNSNNNNNTKN......TH..........C..VR.R.NH.EV	193

TaELF3c	--..T---...S.SQ.RDNNSNN-NNTKN......T...........C..VR.R.NH.EV	109

SbELF3a	--.PT---...S.RQ.E-------NGNKN......A..........LY..V....S..--	191

SbELF3b	--K.I---.G.T.QH.GG------NTI-K.....VV...E.M...IC.PRIYRY....HA	181

SvELF3a	--K.I---..ST.QH.VG------NINKN.....VA.N.E.M...IC.PRFSRY....HA	186

SvELF3b	--.PT---...S.KH.E-------NTTKN......T..........LY..I.......--	193

OsELF3a	--..R---...APQQ.VE------KGIKS...R..AD..E.I......ARF.QY..K.RA	188

OsELF3b	--R.T---..-APQR.AE------NTIKS....R.AD..E.M.....N.RF.QY....NA	192

AtELF3	--------------------------------.MVREEE..A..VYIN.RRSQSHGRTKS	145

BrELF3a	--------------------------------.IAREE...A..VFIN.RR.GRG---KS	126

BrELF3b	--------------------------------.T.REE...A..VYDNDSSTRFQ---SP	142

VvELF3a	--PLQ---.N.SYSQP----------HSL.KL.NHRGE..SE..TFCQ..II.DYYNAPQ	177

VvELF3b	--------------------------------..QGDE...R..IFVH..TEF.-GRNQN	145

PtELF3a	--------------------------------..VGEE......VFVH..KGQGQ.KMQT	146

PtELF3b	------------------------------------------------------------

GmELF3	---------------------------------.KVDE...R..VY.H.RTGQCNDKSVE	142

CpELF3	--------------------------------N.GGDE...R..VFVRF.INQ.HNTIQN	146

Consensus GX-QEKSTPFPATSPXKSX-PXX-K------NTXKRHLEGINVSDVRSR-----------	214

PpELF3b	NPP.VQQSFGGGR.VQ.GKGLVTE.QSPAFSTQLR.ISKEKSM-------------EDDA	253

PpELF3a	VPPNRQGSRQQRAFV------RGKERSSPCM.QVR.L.TEKST-E.V.S-----VRKDDG	241

PpELF3c	S-RE.QLP.GCKNFV-------ME.RTSACV.QVR.IVTEKSI-E.T..-----KRKDDA	247

SmELF3	----------------.PA-AVPARS------SK---Q.HFASAS.APS---------N-	185

BdELF3	RI-.....A..S...Y..G-.TMS.SSAKCS..D..Y...T....M...-----------	225

HvELF3	------------------------------------------------------------

ZmELF3	-----.L.L..T...C..V-.AK------YSS.D..R...MDA...K.K-----------	217

TaELF3a	RI-..NA........Y..G-.TVS.PTAKFP..D..Y...R.A..T...-----------	239

TaELF3b	RI-..N.........Y..G-.TLS.PTAKFP..D..Y...R.A..T..M-----------	240

TaELF3c	RI-..N.........Y..G-.TVS.PTAKFP..D..Y...R.A..T.AM-----------	156

SbELF3a	-----.L.L..T...C..V-.AK------YSSNG.......D....K.K-----------	228

SbELF3b	----D..K.QST.N.H..P-S-MS.SSAKCYS.VNK..DRM.EA.M.LM-----------	224

SvELF3a	.V-.D..N.LS..N.H..P-SAMP.SSAECYSAVN....R.DE..M..M-----------	233

SvELF3b	-----.F....TK..Y..M-.AMS.SSAKCS..D.T....MK...AI.M-----------	236

OsELF3a	.V-..E...LV.L..H..P-.AVS.SPTKCY..VSKN..R......K..-----------	235

OsELF3b	.V-.DQ...LV.AN.H..P-STVS.SSTKCY..VSKK..R.H....K..-----------	239

AtELF3	.IEK..H..MV.P.SHH.I-RFQ--------EVNQTGSKQNVCLATC.KPEVRDQVKANA	196

BrELF3a	.LKRGVGL---.------------------------------------------------	135

BrELF3b	.RRNGEKRTTLF------------------------------------------------	154

VvELF3a	NMNK.RA.C---LDLNS.V-QLE--------SACGK.QRNL..-----------------	208

VvELF3b	SIDR.IL..SSS.CLGH.I-NLQ--------.ACEKE.KQTSSNGLSV.QDKRSQGDVNR	196

PtELF3a	SAA...LASLCP.YLGH.S-RIQ--------.AGDNG--H.GSTGLNL.PDTSNQSEDNL	195

PtELF3b	---------------------------------------------------MRIQIEDNG	9

GmELF3	SFNGK.L..TGSRYFGG.I-SGQ--------SDCE.DPKQFGS.V.NM.KDVRSEIDVLP	193

CpELF3	SVDR..V.HVNV.Y.GS.M-KLQ--------.LSHKEPKQSSS.GLTR-QEVRHQGNEKQ	196

Consensus -DSXSIKE-KEP-------XE-EERSSSFQAXKEKDG-K-DAK-S--------S-SRXDR	252

PpELF3b	SWT-.ASDDCLTVSFIDNDLNTQ.I.N-----RKL..VSR.CDGH-------QTELGRNA	300

PpELF3a	RVP-TASGECAVVSVVDNDVGS..VNP-----S.L..APVACDGG-------FVQPSTHE	288

PpELF3c	SWH-TAS.ECTTVSAIDNDVDSQ.FRH-----T.L.RVPVGHRG--------HVKPSSCK	293

SmELF3	-A.T.AT--.S-----QRQV.VR.T---T..R.D-------SR------------P.-N. 214

BdELF3	-..P...-D.A.LKTMT-NLDV........IS...A.-.A.D.I.----------.HR.K	271

HvELF3	------------------------------------------------------------

ZmELF3	-GPSG...-...VQV-RIDL.DK.TTP...VLND.TW-SP.P.L.----------.HM.. 263

TaELF3a	-..P..IRD.A.ANTTTNFL.T...T....FSA..TMG.R.D.G.--------.Y..VKE	290

TaELF3b	-..P..IRD.A.ANTTTNFL.A...T....FPA..TMG.R.D.G.--------.Y..VKE	291

TaELF3c	-..P..IRD.A.ANTTTNFL.A...T....FSA..TMG.R.D.G.--------.Y..VKE	207

SbELF3a	-GRSG..D-T..VQV-RIDL.D..TTP...IL.D.T.-RP.P.V.----------PFM.. 274

SbELF3b	-N.PKV..-..AVQV-PKGV.VK.KD..I..-S..FKD.-Y..----------LCQMRNK	269

SvELF3a	-S.SK...-..SVQG-SKIV.V..K.LPV..F...FKN.-...----------ACQMG.N	279

SvELF3b	-G.PG...-...TKV-RIDL.I...T....TS...S.-RL.P.V.----------.YR.K	282

OsELF3a	-G.QKD..-TG.AQT-LKNV.V.H-F...E.S.DMF.S.-H..----------VCPK---	277

OsELF3b	-TPLKD..-M.AAQT-SKNV.V.K-....H.S.DMFESR-H..----------VYPKM.K	284

AtELF3	RSGGFV------------------I.LDVSVTE.I.LE.--------------.A.SH.. 224

BrELF3a	--A-------------------------NS.TEDM.LAS--------------TS---.. 151

BrELF3b	--GDQA------------------KG..SSKRHGM.LE.--------------.A.GCE. 180

VvELF3a	--.-HSR.SIS-----------ATA..GLLTRD.IFEPSKR.RT.LTKENKRNVVVDTN. 254

VvELF3b	K.YV.SR.YS.K------------SA.NLSTKEKI.ESMKQVNKPSSEDFRDQ.SANFSS	244

PtELF3a	EVCV.SSDHIAR------------H.TNLRTREKN.IPE-EGNA.QNQQYRNNLV.NFT. 242

PtELF3b	KVFEACQNSM.KFAT-VTSIKDKPSV..SSGKISSTKSLKRTYS.SNQEYRKN.VNVLKC	68

GmELF3	QV.-TS..QASM------------SVR.ISTRENIHTLLRQ..VTPNREFQDCHV.KFN. 240

CpELF3	KNCVLR..HS--------------------.RVKI.EPVKEISG.SRQQLGDGG-----K	231

Consensus LSDINVSDXQHXXXEGHQARSRNXNA-VESQNPPKAGNXSS--P---LXS-N--SXLLEX	303

PpELF3b	VGKKKS--------K.--NKVDGGQIDEDLTK.SDTSAVVI--.PEI.LED-----ESLR	343

PpELF3a	IGQEEL--------LLQ.RDEAR-EKV.RPI.ASETFAAIV--.PVSVHRA-----NSLT	332

PpELF3c	IRQED.--------RI.GNEV.--KSFAGLAKITETHPT-----TARIS.M-----HTLI	333

SmELF3	A-------------P.DLGDQE.GS---------EVRVMA.SS.SHSYE.NE-------D	245

BdELF3	...L..F.K..ART.V....RT.E..-A....A.....G----.SSTNVER.GA.N...K	326

HvELF3	------------------------------------------------------------

ZmELF3	.KK------..AEA.SY.I.T..E..-..T.S...N.VSLLSK.YVDRREQ.GD.D..GH	316

TaELF3a	T.S.....K..SRN......T..E..-A....AA....G----.YSTDVAC.GA.N.S.K	345

TaELF3b	T.S.....K..SRN......T..E..-A....A.....G----.YSTDIAC.GA.K.S.K	346

TaELF3c	T.S.....K..SRN......T..E..-A....A.....G----.YSTDVAC.GA.N.S.K	262

SbELF3a	.KKY..A.K.YSEA.SY.M.T..ED.-.KT.....N.SVLLSK.YDD-REQ.GD.DI.KH	332

SbELF3b	V.N..R..N-----NSC.PT.V.GKS-T.AK..TATR.P..CK.CTDVD.S.WN.N...R	323

SvELF3a	ATN.DSY.NP.FGNSRR.PT.M.GSS-M.AK..TTTR.TV.CK.CTD.NDS.KN.N..DR	338

SvELF3b	.NKY..A.K.SSEIASY.T.N.KE..-G.T....E.EMAP.AK.YAG-MEQ.GN.D...L	340

OsELF3a	TGT..DL.EP.LENSE...T...GSS-.KF....VRR.TI.AK.SPGIENT.GHCN.PQG	336

OsELF3b	TGI..D..EP.GGNS....T...GGS-MKF....MRR.EI.SN.S--SENTDRHYN.PQG	341

AtELF3	VN.Y.A.LR.ESRNRLYRDGG------KTRLKDTDN.AE.H------.AT----ENHSQE	268

BrELF3a	VD.GRINN----------------------------.VE.H------MEA----.E--.E	171

BrELF3b	VNASFLLRQE--------------------------STS.R------.EL----D-----	199

VvELF3a	.E..HD.LH.GSMTLPENTDLP.PS------KAVGG.YV.T------SPKP-----S.LD	297

VvELF3b	.H.TDGCLL.GNRAGAQLGEPDHVDCVSVET-ARD.W.A.R------VR.SS----YSGD	293

PtELF3a	.HENDTCLQ.ETSARLQSNH.EHGDHVP..R-RQ.EKINIF------QPGND---SH.RK	292

PtELF3b	.PGT.EQLN.ELVTMPDKTVIGDNI-L..YRVVTGKE.P.K------VR.ELYSRA..QD	121

GmELF3	.QQGETCLQLECGV.SRSNDIGDNGCL...ARETDK..APT------------------A	282

CpELF3	.N.DVAQWQ.GARTGLQRVD.GPVDDGDSLR-DVDK.IMPQ------.R.NS---CSRDD	281

Consensus GLRE-G-KRKRSX-----------------HNVEQNDDLS-DSSVESLPGLEISPDDVVG	343

PpELF3b	VVP.QDIFSSQ.----TENRSREVSETSQRSR--GGSAETES.IL.D..LGSSCFQELLD	397

PpELF3a	.AS.RNDFSNQT----SENQRGEGSKPSRLRR--DSSMETGS.ML.NVVMGS.EFR...D	386

PpELF3c	APS.QDIISNI.----SENQSEEISELYQQKR--GSSARTGSLML.NV.TDFVQLR...N	387

SmELF3	NG.---IDNQSDEELEGRC---------SPEE..AS..SATSVVENNAAPSA.TSKEIMS	293

BdELF3	...VT.E.....EG----------------.HNV.K..S.-.L................. 369

HvELF3	------------------------------------------------------------

ZmELF3	....T.V.....H-----------------.D........-.........M........S	358

TaELF3a	....T.E.....TG----------------.HDV.R..S.-........D.......... 388

TaELF3b	....T.E.....TG----------------.HDV.R..S.-........D.......... 389

TaELF3c	....T.E.....TG----------------.HDA.R..S.-........D.......... 305

SbELF3a	...DT.E.....H-----------------.G........-.....F.A.M......... 374

SbELF3b	SP..G.A....QH-----------------.SG.......-.D...CI..W.L...EIA. 365

SvELF3a	S...A.S....GH-----------------QD........-.....CI..W.V...EI.. 380

SvELF3b	....T.E.....H-----------------.G..H.....-........EM........S	382

OsELF3a	..K.A.T....LEAQD---------------.A.KI....-.....CITAW.....EI.. 380

OsELF3b	.IE.T.T....LLEQHDAEKSDDVSRLLEQ.DA.NI..V.-.....CIT.W.....KI.. 400

AtELF3	.HG-----SPEDIDNDREYSKSRACAS**L**QQI.E.AS..V.D..M.D.ISSIDV....... 323

BrELF3a	.HG-----NLND---EYCTSGGGGYTSLQQEINNEEEASDDN.M.D.VSS.DV...E... 223

BrELF3b	QDG-----DETG---VMETD--DGVE----SHGDP..VDND.V.DD.ISSVDV...E... 245

VvELF3a	.H.-----.SG--GLEHELEEQKESESLQMGSIDR.CHV.NANM.G.ML.FD........ 350

VvELF3b	..G-----SPSEPDNDGACRGDKTCGTLQKG..DT.....ET.M.D.MS..D.T...... 348

PtELF3a	DCS-----SPNEPEIDSECFGDKTCGSLQFR.GDKSN.A.ET.M.D.VSV.D........ 347

PtELF3b	DN.-----NRCGLEKRSKYREDKQSGSLKAGDL.R...AAET.M.D.VTA...T...... 176

GmELF3	NQT-----SPADAINDTEHHDTRMGSPIQRG.LNES.NA.KI.M..N.STVR........ 337

CpELF3	PSS-----SPDEHDNDSGCTRNNARGSFQLV.GDKS..V.ET.M.D.ISS.D........ 336

Consensus AIGQKHFWKARRAIVNQQRVFAVQVFELHRLIKVQKLIAASPHLLIEG------DPCLGK	397

PpELF3b	HF..QEL...QK..IR..K..SR.......V.E..R.L.KL.S.SFRLDEV--IEAEGDN	455

PpELF3a	.F..QE...VQMM.IR..KL..L.......A.E..Q.L.KALCFSLHVDE---VPQKVDN	443

PpELF3c	IF..QG...TQKV.IR...I.SK.L.D...IME..H.L.KELCFS.TVDEDK-VEEEDDN	446

SmELF3	.V.DEE...M.K.MQR..TI.QK.L......T...H.M.N.KISDPSKTQDEKT.KRV.S	353

BdELF3	...P............................................------.....S	423

HvELF3	------------------------------------------------------------

ZmELF3	...P.......................................V....------...... 412

TaELF3a	...P............................................------.....S	442

TaELF3b	...P...........................................W------.....S	443

TaELF3c	...P............................................------.....S	359

SbELF3a	...P............................................------...... 428

SbELF3b	...P..........Q..............K..................------..V..N	419

SvELF3a	...P.......K..Q..............K..................------..V..S	434

SvELF3b	...P............................................------....D. 436

OsELF3a	...A..........I.......A......K.V...........V....------.....N	434

OsELF3b	...T..........M..............K.V...........V...S------.....N	454

AtELF3	IL...R..R..K..A.........L.................D..LDE------ISF... 377

BrELF3a	VL...R..R..K..A.........L..........R......DV.VDD------MSYV.. 277

BrELF3b	V....R..R..K..A....I....L..........R...S.SDV.LDE------ISY..N	299

VvELF3a	I..H.....T.N.......................R...G..EGQLDN------NLY.AE	404

VvELF3b	I..................................R...G....MVDE------SAY... 402

PtELF3a	I.............A......S..L..........Q...G...V.L.E------EVH.A. 401

PtELF3b	V..E.Q.....T..........A................G.....L.D------NLYV.R	230

GmELF3	I.............A....................Q...G..DI.L.D------GAF... 391

CpELF3	I.............I........................G...I.L.D------SAF..V	390

Consensus ALX---------------LXSENVEKQPLSA---K--NKDD--E-TTLQQXECSKENTEG	434

PpELF3b	VPL------LDSPVAEVPEV---------------------------PEVP.V.GRSL.K	482

PpELF3a	I.R------PSSPVAEVPETP.AL.IPKA-----------T----EIPEVP.V.ERRF.T	482

PpELF3c	VSV------PSSPLAVVPEVH.DS.ISGA-----------S----QVPEIS.TPDRCLDT	485

SmELF3	EPT------------------HCG.APKT.----.PGP.P.----------------QAP	375

BdELF3	..ATSKKK----------.AAG......P..---.--.--KDDAQL....V.Y..D.I.. 466

HvELF3	------------------------------------------------------------

ZmELF3	S.AVSKKR----------.AG-D..T.LE..---.--.D.G-VRP.---.L.H...K..A	452

TaELF3a	..VTSKKK----------TAAA.....L...---.--S...DDAQL....A.Y..D.... 487

TaELF3b	..VTSKKK----------TAAA.....L...---.--S...DDAQL....A.Y..D.... 488

TaELF3c	..VTSKKK----------TAAA.....L...---.--S...DDAQL....A.Y..D.... 404

SbELF3a	S.AASKKK----------.AG-D....LQ..---.--.N.E-VQP.QQ..L.H......A	471

SbELF3b	..MGKRNK----------.PKG.LKV.T..I---T--....--IQP..E.P.L..Q.... 462

SvELF3a	..VGKKTK----------.PKG.LKV.T..I---A--....--IQP.PE.P.L..Q.... 477

SvELF3b	..AASKKK----------.AGGDA...HQ..---.--Y...-VQQ.-...L.H..D...A	479

OsELF3a	..LASKKK----------MAE..LKA..VLV---A--TN..--VQPS..EP.L....S.E	477

OsELF3b	..LGSKNK----------.VE..LKA...LV---A--TI..--VEPS...P.V......D	497

AtELF3	VSA------KSYPVKK-L.P..FLV.P-PLPHVVVKQRG-.-S.K.DQHKM.S.A..VV. 427

BrELF3a	VAA------KSYPVKKLH.P.GFLV.P-PLPQVI.HRSS.Y-S.K.DQHKM...A..VV. 329

BrELF3b	VPV------KK------L.P..FIV.P.PLPHAT.HRRG-.-P.K.DQDKM...A..VV. 345

VvELF3a	PASSTKKLPSESPTKK--.P..S.QQL.SQI--AD--P.NG-SQ-KQDSNN.GAVKDAV. 456

VvELF3b	PSL------KSSPAKK--.PL.Y.V.P.PNM--VM--H...-Y.-RASH.L...A..AV. 448

PtELF3a	PPM------KGSPCKN--.P..CAVTP.VHV--A.--H..N-S.-NPNHKM..FA..AV. 447

PtELF3b	.SL------KVSQINK--VP.KCAMVD-------.--P..H-SQ-KQHTSADFAG..VV. 271

GmELF3	SPP------KGSTPKK--.AL.Y.V.PRQQN--L.--R...-S.-KLNHKM...A..AV. 437

CpELF3	PSL------KSSSAKK--FT..Y.V.L..Q.--VN--C...-S.-KANDKM...A..AVV	436

Consensus NQPSPSQDD-----QHXNQAAXNGAFSSNPPXMPAXPD---NKQNNWC--P-PP-----Q	478

PpELF3b	DISR..DEI---------SSQVPESL.QPAYPLSTP.QQGF.PASP.QAA------TYSS	527

PpELF3a	KVCKDLNEN---------SRSVPESV.QPLYPL.LPFQQGF.SGSA.PAG------PYAA	527

PpELF3c	.VCEA.NEA---------PRPVPPP.PAPMYSL.VLSQHSFTPESL.QAP------PYAT	530

SmELF3	APKT...TQ--------PLSTTPY.YTTH.RPIQ.--A---.TI.YAT--SH.A-----Y	415

BdELF3	..A......-VVVV..N....S..GDT....AI..A..---...S...--TP-.-----. 514

HvELF3	--------------------------------...PT.---.......--AP..-----. 18

ZmELF3	........E-QAATNG-----------DVAAS.HTPS.---...KS..--IPA.-----P	490

TaELF3a	..A.....NDVVEVR.E....S...V.....A...P..---.......--AP..-----. 537

TaELF3b	..A.....NDVVEVR.E....S...V.....A...P..---.......--AP..-----. 538

TaELF3c	..A.R...NDVVEVR.E....S...V.....A...P..---.......--AP..-----. 454

SbELF3a	.........-AAGV..N....I...V.....S..TPS.---....S..--IP..-----P	520

SbELF3b	.PSHH.R..-GLDDN.HD...A.ET.T....AI.VA..---.......--MN..-----. 511

SvELF3a	.P...CR..-GLGGNGHD...T.ET.T....V...A..---.......--MN..-----. 526

SvELF3b	A....T...-VVAA..N....ATA.VN....T..TPS.---....S..--IP..-----P	528

OsELF3a	.P...-R.T-APVSG.HD.T.KI..SK..LRAT.VAS.---.R...CGVQLQ..-----. 527

OsELF3b	SP...-H.T-GLGSGQRD...T..VSK..RRAT.VAS.---......GVQLQ..-----. 547

AtELF3	RLSNQGHH------.QS.YM----P.AN...AS.--------AP.GY.FP.Q.-PPSGNH	468

BrELF3a	RL.NQG.GHHH--Q.PS.YM---MP.AT.Q-P----------NA.GCYYP.Q..TPSGGN	373

BrELF3b	KLSNQG.QH-----.PS.YM----P.A....T----------AV.GCYYP.---PPSGGN	383

VvELF3a	.P.FA.PV.V---DKIYAAQQT.QRPLE.Q.PVSMAT.---TQPTS..FH.-SQ-----G	504

VvELF3b	KTHL..VKNG---NPPS.Y--GP--YIG...PA..PT.---S.MGP..YPQ-..-----G	492

PtELF3a	KT.FA.VKNG---QPP-.F--GPH---AG.TTV.MAS.---T.MAP..FH.-..-----G	489

PtELF3b	KL.L..TN.E---TSKEPI-SQRSNY.GSA.PA.VATT---A.PSP..YP.--.-----G	317

GmELF3	KTSLS.VK.G---SHLSKC--TP--.PG.QHQTNVAA.---SGMGP..FNQS..-----G	482

CpELF3	KTSL..ARNG---S.PS.Y--S.QQCLR..LPA.VA..---A.VGP.SYH.-S.-----G	482

Consensus NQWLVPVMSPSEGLVYKPYPGPCPPA------GSFLA-PFYG-SCGPLSLPSTA-GDFMN	529

PpELF3b	.PYAARMS..---HM.V......S.GYGVYPMMGAPM-SM..--------NFGGMQASRF	575

PpELF3a	.PCATRM...---YM.V..S.....GYGAYPMMDAPM-.RF.--------YPGVMQTVRF	575

PpELF3c	.P.APRMA..---Y..V...PT.S.GYGAYPMMGASV-.V..--------HPG.MPV.RF	578

SmELF3	...YA.LPFQ------F.FQQ.F.VTAQVFNTPYYP.-.L..GGAA.PF..PPDPRQ..S	468

BdELF3	...................T......------.....-...A-..A........-..... 565

HvELF3	...................T......------.....-...A-..A........-.E... 69

ZmELF3	S...I..............T.H...V------..L..P..FA-.YPTS.SSTAG-....S	542

TaELF3a	.............F.....T......------..I..-...A-..A........-.E... 588

TaELF3b	.............F.....T......------..I..-...A-..A........-.E... 589

TaELF3c	.............F.....T......------..I..-...A-..A........-.E... 505

SbELF3a	S..................S.H....------...M.P..FA-....V......-..... 572

SbELF3b	...................A.....V------.NL.T-...A-N.T..R....-------	556

SvELF3a	...................A.....V------..L..-...A-N.T..K....-------	571

SvELF3b	...................A.H....------.....-...P-..A.V......-....S	579

OsELF3a	....I..............S......------..I..-...A-N.T..R....T-..... 578

OsELF3b	..........L........S......------..I..-...A-N.T........-..... 598

AtELF3	Q...I.........I...H..MAH-----------T--GH..GYY.HYMPTPMVM-----	510

BrELF3a	Q...I.........I...H...G.---------GHTG-.VC.GYY.HFMPAPMFM.G---	420

BrELF3b	Q...I.........I...H...G.--------------.VC.GYY.HFMPAPMMM.S..-	428

VvELF3a	..............I....A.Q.S.T------PRLMS-....-NYR.VN.TVMG-...F. 555

VvELF3b	H.....................G-----------.MS-TVC.-G...MGSAPMT-.S.I. 538

PtELF3a	L............F.....TA.G-----------.MG-SGC.-G...FGPIPLT-DN..T	535

PtELF3b	...................A.....V------SR.ME-.V..-....I..APGG-...L. 368

GmELF3	HP..I...T.............G-----------.TG-TGC.GG...FVPALLG-.S... 529

CpELF3	H.....................G-----------.AG-SVC.-G...FGPTPMT-.A..A	528

Consensus SXYGIPMPHQPQXMGVPGPP--------------XPMXPGYFPPFSMP--VMN-PVVSXS	572

PpELF3b	PTWQQQGMS..WGPSD.AAAGAVWYGQQMVPAVGPATNM.VI.MVT----NCERQTS.GA	631

PpELF3a	PTWSQ.SMP.TWTSLNAAAAAAAWYGQHVSA-AVPTANT.VAS.AR----NSQRLIS-SG	629

PpELF3c	PAWPL.GMSH.WNARDSV-EAAAWYGQQAVPGAPPVPSK.FSS.AC----N.EHFAS.SG	633

SmELF3	PPIQLQPWP.HPGSLYSD.AALQWMGFVN-----P.GGAT.R..AAES--SLQ-----A. 516

BdELF3	.P...H......H..LG...---------------..P.M........--...-....A. 607

HvELF3	.P..........H...G...---------------A.P.M......V.--...-....S. 111

ZmELF3	.AC.ARL----------------------------MSA.V...S....--A-----..G. 567

TaELF3a	.P..........H...G...---------------A.P.M......V.--...-....S. 630

TaELF3b	.P..........H...G...---------------A.P.M......V.--...-....S. 631

TaELF3c	.P.........HH...G...---------------A.P.M......V.--...-....S. 547

SbELF3a	.A..VA......H.......---------------..P.M........--...-.A..A. 614

SbELF3b	-P..V.......Q.VP..A.---------------A.HMN........--...-.GTPA. 597

SvELF3a	-P..V.......H.AP..A.---------------A.HMN......V.--...-.GAPA. 612

SvELF3b	.P..........H.......--------------P..P.M........--...-TA..A. 622

OsELF3a	.A..V.I.....H..A..T.---------------T.PMN......V.--...-..ALA. 620

OsELF3b	.A..V.......H..A....---------------S.PMN......I.--...-.TAPAP	640

AtELF3	-------.QYHPG..F------------------P.PGN.....YG.MPTI..-.YC.SQ	544

BrELF3a	---.GQP.PFHPG..F------------------PSHGN.....YGGI--M..-.YY.GQ	456

BrELF3b	---.GGP.PFHPGV-----------------------GN.....YG----M..-.YG.GH	457

VvELF3a	TT....IS-LQ.GIEIVPSN--------------PFLGQT....YGK.--LV.-.ST.G. 597

VvELF3b	PA..V.SS.HH.GI..HPGT--------------P.IGH.....YG.S--...H.TI.G. 582

PtELF3a	.A.A..TS.YH.GI..SPGA--------------P.VGNAC.A.YG..--G..-.AI.G. 578

PtELF3b	AA.SVSAS-NHEEI.ILPGN--------------PHFGQTF.Q..G..--...-.SICD. 410

GmELF3	PG....TS--H.GV...PDT--------------H.GSH..L..YG..--...-SSM.E. 570

CpELF3	PA..V.G.-PH.GVA.LPGA--------------P.VGHS...SYG..--I.S-.SF.G. 570

Consensus A------VEQVSHVAAPXPNGH--------------E-HSRSSCNMSNP---VXXSGVWK	608

PpELF3b	.---QLIQRNYAQKGSARQA.EKS----------------DVGG.R.KLVNA.ESC.WA. 672

PpELF3a	S---GQS..NTQQAGVGS----------------------DVGV.LV.KVGPLDDR.WA. 664

PpELF3c	S---RQS..RTAQMGNDRSA.KMS----------------DTGA.R..EGDF.EGCDRA. 674

SmELF3	S-------------------I.DD--------------.QSRKDRGIHLRSQGSS..FSL	543

BdELF3	.------.....RI.PAR..A.-------------V.HY..N....R.EA--MS-A.I.R	645

HvELF3	.------.....R...AR..T.-------------L.H........R.EA--.SVG...R	150

ZmELF3	.------.........SQ--------------------.K.N..SEAV------------	589

TaELF3a	.------.....R..TAR..T.-------------V.H........R.EA--.SAG...R	669

TaELF3b	.------.....R...AR..T.-------------V.H........R.EA--.SAG...R	670

TaELF3c	.------.....R...AR..T.-------------I.H....------------------	588

SbELF3a	.------.........SQR...-------------I.Q.T.N...A.HLRSEAVSA...R	655

SbELF3b	.------...G..A...Q.Q..-------------M.QQ.LI.....H.------..I.R	632

SvELF3a	.------...G..A.V.Q.H.R-------------A.QQ.LI.....H.------..I.R	647

SvELF3b	.------.........SR....-------------I.Q..........LRSEALSADI.R	663

OsELF3a	.------...GR.PSM.Q.Y.N-------------L.Q...M.....H.------..I.R	655

OsELF3b	V------...GR.PSM.Q.Y.N-------------F.QQ.WI.....H.------..I.R	675

AtELF3	QQQQQQPN..MNQFGH.GNLQNTQ-------------QQQQR.D.EPA.QQQQQP---T. 588

BrELF3a	QQQQQQPN..MNNN-----IQ------------------QQ..V.EATSQQQQQP---T. 490

BrELF3b	QQQ-QQPS..MNQFVH.VNTQ------------------QQ..V.EAIS--QQQP---T. 493

VvELF3a	.------...MNLSVGDQSSRPNNHVSVGDINATVQSV.YQ.LG..PSQKSGGILGCFG. 651

VvELF3b	.------...MNRF.GHGSLSQSGQLSGGGA-S--FNMQHQN...VPT.K-RAIPQ..-. 631

PtELF3a	--------------.GSGSC.QTAQFPGGIL-S--SNMPHQ....ERTQKSEAVLE.M-. 620

PtELF3b	.------...IRPRIG.QS--KDNQLAVGDV-N--FNIPLQ.......QMSR.ISCC.EN	459

GmELF3	V------...GNQFS.LGSH..NGHLPGGGKAN--HNTNNK....LPVQRNGAISHVL-. 621

CpELF3	.------...MNPFST.GGH..IANLSGGGA-K-------Q....L.TQKTGTIPQV.-. 615

Consensus FHASRDSELQGSSA-SSPFDRQQG-G------GRGPAPPFPAAPA---------------	645

PpELF3b	GWG.--T.DRR------------.HREPDASRN----KE.G.V.LGDE-----SERQDGE	709

PpELF3a	GWG.--..R.R------------AS.-EGNPQN----KTSF.GLRGDN-----GNCQNSE	700

PpELF3c	VGGG--G.-VR------------VR.DRNSAQK----TGDI.DLRGEE-----TDCEDGA	710

SmELF3	IQ-RA.--------------------------SSS...APSPS..AE------NRRQQHQ	570

BdELF3	..........A...A.........E-------A....A.P.---IP-TSSAGNG------	688

HvELF3	..S..G.K......A.........Q.E-----A..H.AAA....PPTSSSAGNGNGNAAQ	205

ZmELF3	-L......V.....-...AS----------------------------------------	607

TaELF3a	..S..G........A.........Q.E-----A....AAA....LPTSS---AGNGNAAQ	721

TaELF3b	..S..G........A.........Q.E-----A....AAA....LPTSS---AGNGNAAQ	722

TaELF3c	------------------------------------------------------------	648

SbELF3a	V.............-.........E.R-------.....-----FPA-SS--VGNRQAQA	699

SbELF3b	.L......P.A...-......LEAQ.D-----.S..VSF..K.SVL------------NA	674

SvELF3a	......R.P.A...-......I.VQ.D-----.S.LVSV..S...Q------------NA	689

SvELF3b	....K.........-..T......E.R-------...Q.-----FPS-SS--VGNGQP--	705

OsELF3a	........A.A...-......L.C--G-----.S..VSA..T.S.Q------------NT	695

OsELF3b	........A.A...-......F.C--S-----.S..VSA..TVS.Q------------NN	715

AtELF3	SYPRARKSR...TG-...SGP.-.IS------.SKSFR..A.VDEDSNINN------APE	634

BrELF3a	SYPRAKKSR.EGIS-----------.------KKKSFQ..S.VDDDDDDNDKINNAAPPT	533

BrELF3b	SYPRARKSR.R.TG-...RGPE-.IS------DTNSFR..SVVDDDDNNE--------PE	537

VvELF3a	.Q..KS....V.T.-.T.SK.A----------QADAL.L..V..K---------------	685

VvELF3b	.PM.K...F...T.-...SE.E.QV.TGDTAE..D.L.L..M...---------------	675

PtELF3a	LR..KNTSV...TG-...SG.V..V.TVQAAD..AAF....VT.P---------------	664

PtELF3b	.QGLKE..I.....-G.LSKMP----------KANAL.L..ME.T---------------	493

GmELF3	HQT.K.F...ET..-...SEMA..LSTGQVAE..DVL.L..MV..---------------	665

CpELF3	HQP.K.TQ....T.-...GE.V-QD.IGNIGE..NAL.L..T..D---------------	658

Consensus QPQ---------------------------------------------------------	648

PpELF3b	TRRVETGGGVGQPNDEGVRMVEREKNSGVLSAHGREVSTRG-GELCDH-GEELGGLELER	767

PpELF3a	DHRYGETER--DVSGCGVRQRTNEDDR---------------GVECDVDSREYREPDSEK	743

PpELF3c	AC.DAEAGQ--ENSDCEVKNHSNERNK---------------GNSKEGAE----------	743

SmELF3	E.R--------------------------QKLVGRF---------CGE------------	583

BdELF3	...PSTGSK--------------EN-----------------------------------	699

HvELF3	...VSSGSQ--------------ENPVA--------------------------------	219

ZmELF3	------SET--------------AA-----------------------------------	612

TaELF3a	...VSSSSQ--------------ENPVA--------------------------------	735

TaELF3b	...VSSGSQ--------------ENPVA--------------------------------	736

TaELF3c	------------------------------------------------------------	708

SbELF3a	.A.ASSGSR--------------EN-----------------------------------	710

SbELF3b	...PSSGGR--------------DQQNHVIRVVPRNAQTASVPNAQPQ------------	708

SvELF3a	...PSSGSR--------------DQQNHVI------------------------------	705

SvELF3b	--.PSSGSR--------------EN-----------------------------------	714

OsELF3a	...PSSGSR--------------DNQTNVIRVIPHN------------------------	717

OsELF3b	...PSYSSR--------------DNQTNVIKVVPHN------------------------	737

AtELF3	.T----------------------------------------------------------	636

BrELF3a	EE----------------------------------------------------------	535

BrELF3b	.M----------------------------------------------------------	539

VvELF3a	------------------------------------------------------------	685

VvELF3b	------------------------------------------------------------	675

PtELF3a	------------------------------------------------------------	664

PtELF3b	------------------------------------------------------------	493

GmELF3	------------------------------------------------------------	665

CpELF3	------------------------------------------------------------	658

Consensus ---------------------------------------------XTRVIKVVPHNARTA	663

PpELF3b	NLPAGARHKEPLEVRTLKRPRTEFPKSDGFPWFPIISPAKRVMQQC-G......RAVSAT	826

PpELF3a	SWPAETSHAGPLQGRSLKHMRSESLEPSASRWFPTLSPAKRVMHKYGG......RAVSAT	803

PpELF3c	LESMGRQQKGSSNGKSLKRMRSESLDPDACRWFPILSPAKVSKHQI-G...A..RAVSAT	802

SmELF3	KQSGGESEH-GVSPYSRPSGFQ------SVKNSSSSSKEQRSSQGSHKA...T.RA.PVT	636

BdELF3	---------------------------------------------PAG..R....T.... 714

HvELF3	------------------------------------------AAAAA...R....T.... 237

ZmELF3	---------------------------------------------QP...R....T.... 627

TaELF3a	------------------------------------------A--AA...R....T.... 751

TaELF3b	------------------------------------------A--AA...R....T.... 752

TaELF3c	------------------------------------------------------------	768

SbELF3a	---------------------------------------------PS...R....T.... 725

SbELF3b	PSSGGRDQQ-NHVIRVVPRNAQT-----ASVANAQTQPSSGGQDQWNH..R......Q.. 762

SvELF3a	--------------------------------------------------R......Q.. 715

SvELF3b	---------------------------------------------PG...R....TS... 729

OsELF3a	-------------------------------------------------------.SQ.. 722

OsELF3b	-------------------------------------------------------S-... 741

AtELF3	-------------------MTTT-----TTTTRTTVTQTTRDGGGV...........KL. 672

BrELF3a	-------------------MMTT-----TTTTTTTVTQTTRDGAGV...........NL. 571

BrELF3b	-------------------MTTT-----TTTTRTTVTQTTRDGGAV...........KL. 575

VvELF3a	-----------------------------IQES-DQLDQIHGNEKQ..A......KHKS. 715

VvELF3b	----------------------------AIPAG---DPQPNGTDQP....R.....P.S. 704

PtELF3a	-----------------------------CPEG---APQHQETDQLSK........G.S. 692

PtELF3b	-----------------------------LQAS---YPNAQTNEQQA.........R.S. 521

GmELF3	-----------------------------EPES---VPQSLETGQH..........R.S. 693

CpELF3	----------------------------AV-------MQPIDTEQQ..........P.S. 683

Consensus SESAARIFRSIQMERKQNDP**XXX*	689

PpELF3b	Q....S.LL...K..QR*--------	844

PpELF3a	Q..T.S.LL...T..RR*--------	821

PpELF3c	L....N.LL...T.KWR*--------	820

SmELF3	A....E.LH...K..PS*--------	654

BdELF3	...............Q.....-----	735

HvELF3	...............Q..G..-----	258

ZmELF3	.....................-----	648

TaELF3a	...............Q..G..-----	772

TaELF3b	...............Q..G..-----	773

TaELF3c	--------------------------	570

SbELF3a	........Q............-----	746

SbELF3b	...................S.-----	783

SvELF3a	...............R.....-----	736

SvELF3b	........E..K...Q...*------	749

OsELF3a	...............Q.D.S.-----	743

OsELF3b	...............QRD.*------	761

AtELF3	..N.....Q...E...RY.SSKP---	695

BrELF3a	..N.........E....YY..-----	592

BrELF3b	..N.........E...HY.SFSNHS. 601

VvELF3a	............E..N.RSTQGDNY. 741

VvELF3b	T.......Q...D....L.ST.----	726

PtELF3a	T..V....Q...EG...Y.SL.----	714

PtELF3b	T.......Q...E....Y.*------	541

GmELF3	TA......Q...EG.....SV.----	715

CpELF3	...V....Q...E...HL*-------	702
